# PAIR-Scan: Single-step identification of highly potent TCR–neoantigen pairs through library-on-library screening

**DOI:** 10.64898/2026.09.07.749948

**Authors:** Marius Messemaker, Connor A. Richterich, Jule Wenner, Yiting Ma, Živa Moravec, Rhianne Voogd, Maike Mussmann, Sam de Paauw, Bjørn P.Y. Kwee, Jeroen Geerligs, Ben Morris, Yaël Winkler, Babet O. Springer, Floris T.B.T. van den Brekel, Jos Urbanus, Aurélie Guislain, Nuno Alfaiate, Frank van Diepen, Martijn van Baalen, Benoit P. Nicolet, John B.A.G. Haanen, Catherine J. Wu, Jani Huuhtanen, Wouter Scheper, Ton N. Schumacher

**Author notes:** These authors contributed equally (Marius Messemaker, Connor A. Richterich).

## Abstract

Tumors contain a mixture of T cells with bystander reactivities and reactivities towards different, frequently patient-specific, cancer (neo)antigens. The one-step identification of highly active TCR–antigen pairs in human tumors would be valuable, both as a monitoring tool and to boost T cell reactivities of interest. Here, we develop PAIR-Scan, an HLA-agnostic library-on-library screening technology that identifies functionally active TCR–neoantigen pairs among tens of thousands of candidate pairs in a single step. We demonstrate the value of PAIR-Scan on a range of tumor samples and for the direct identification of TCR-recognized minimal peptides. In addition, we demonstrate that PAIR-Scan correctly ranks TCRs reactive to the same antigen by their relative tumor-killing efficiency. Together, these data demonstrate the value of PAIR-Scan for both the dissection of T cell responses in clinical samples and to generate large-scale datasets for the development of predictive models of TCR reactivity.

## Main

Neoantigens (neoAgs) that arise from nonsynonymous tumor mutations in expressed genes are a frequent target of tumor-reactive T cell responses^1^. The clinical relevance of neoAg-specific T cell reactivity is supported by the relationship between tumor mutational burden (TMB) and immune checkpoint blockade (ICB) response rate in both metastatic disease and in the neoadjuvant setting^1–3^. However, our understanding of neoAg-specific T cell responses in individual patients is complicated by several factors. First, only a small fraction of expressed mutations leads to immunogenic neoAgs, and the large majority of these are ‘passenger’ mutations that do not recur across patients^1^. Second, while tumor antigen-specific T cells show increased expression of T cell activation- and T cell dysfunction-related signatures^4^, transcriptional information on tumor-infiltrating T cells is insufficient to identify tumor-reactive T cells with high precision. Third, computational models of T cell receptor (TCR)–Ag pair reactivity at present only have a limited ability to correctly predict recognition of patient-specific, and hence ‘unseen’, epitopes^5^. For these reasons, analysis of neoAg-specific T cell immunity in patient cohorts requires scalable technologies to measure TCR-(neo)Antigen interactions on a patient-specific basis. Identification of reactive TCR–neoAg pairs using such technology may serve at least three purposes. First, it may be used to monitor immune activity before or during therapy and to guide patient selection, e.g., identifying those with a broad pre-existing tumor-reactive T cell response that may benefit from ICB. Second, it can enable therapeutic strategies that selectively expand the tumor-reactive T cell pool, either through vaccination^6^ or adoptive cell therapy^7^. Third, it contributes to fundamental insights into cancer-specific T cell immunity, such as the phenotype of T cells reactive to (specific classes of) tumor antigens, and by generating data sets that enable improved *in silico* prediction of TCR–Ag pair reactivity. However, not every identified TCR–neoAg pair should be considered of equal significance for these purposes. Specifically, TCRs can vary widely in their capacity to induce tumor killing, and this variety in cytolytic potential is also observed for different TCRs that are reactive to the same tumor antigen^8–11^. Hence, assays that do not just detect the mere presence of interacting TCR–antigen pairs but also rank such reactive pairs by potency are preferred.

We set out to develop a screening technology that can identify reactive TCR–neoAg pairs in a single step and also rank identified reactive TCRs by their potency. In order to avoid the need for viable tumor-infiltrating lymphocytes (TIL) and tumor cells, we focused on approaches in which genetic information on tumor material (i.e., mutations in expressed genes and intratumoral TCRαβ pairs) is used to create synthetic antigen libraries and synthetic TCRαβ libraries that cover this combinatorial space. Introduction of the resulting antigen libraries into autologous (patient-derived) B cells allows interrogation of antigen presentation by any of the patient’s human leukocyte antigen (HLA) class I or II alleles^12^. Furthermore, introduction of the resulting TCR libraries into reporter T cells allows analysis of antigen recognition without variability in TIL frequency or TIL dysfunction as a potential confounder^13^.

## Results

### Interrogating the intratumoral TCR–neoantigen space using T cell–target cell doublets

To estimate the number of candidate TCR–neoAg pairs that should be interrogated in a single screen to cover the combinatorial space in most human tumors, we determined the number of unique TIL TCRαβ amino acid clonotypes and the number of expressed non-synonymous single-nucleotide-variants (SNVs) and insertion-deletions (indels) across a range of tumor samples. In 95% of the sampled tumors in recent single-cell RNA & TCRαβ-sequencing studies (listed in Supplementary Table 1), 800 or fewer CD8^+^ and 2,552 or fewer CD4^+^ clonotypes were detected (Fig. 1a and Extended Data Fig. 1a-c). Across cancer types, the median number of CD8^+^ TCRαβ clonotypes was correlated with median intratumoral CD8^+^ T cell abundance, as determined by whole slide imaging^14^ (ρ = 0.59, Extended Data Fig. 1d). Prior work has shown that tumor-reactive TCR clonotypes are enriched in T cell populations that are clonally expanded or that display a dysfunctional or tumor reactivity signature^4,15,16^. When restricting to TCR clonotypes expressed by at least two T cells, 213 or fewer CD8^+^ and 345 or fewer CD4^+^ clonotypes were detected in 95% of tumor samples (Extended Data Fig. 1e,f). Likewise, examining tumor mutational burden in the 95^th^ percentile, at most 365 candidate neoAgs, derived from either SNV or indel events, were detected per tumor sample in the cancer genome atlas (TCGA) dataset (Fig. 1b). Together, these data indicate that a screening approach capable of testing several hundreds of candidate TCRs against several hundreds of candidate neoAgs is required to cover the combinatorial TCR–neoAg space that is routinely sampled in most human tumors.

**Figure 1:**
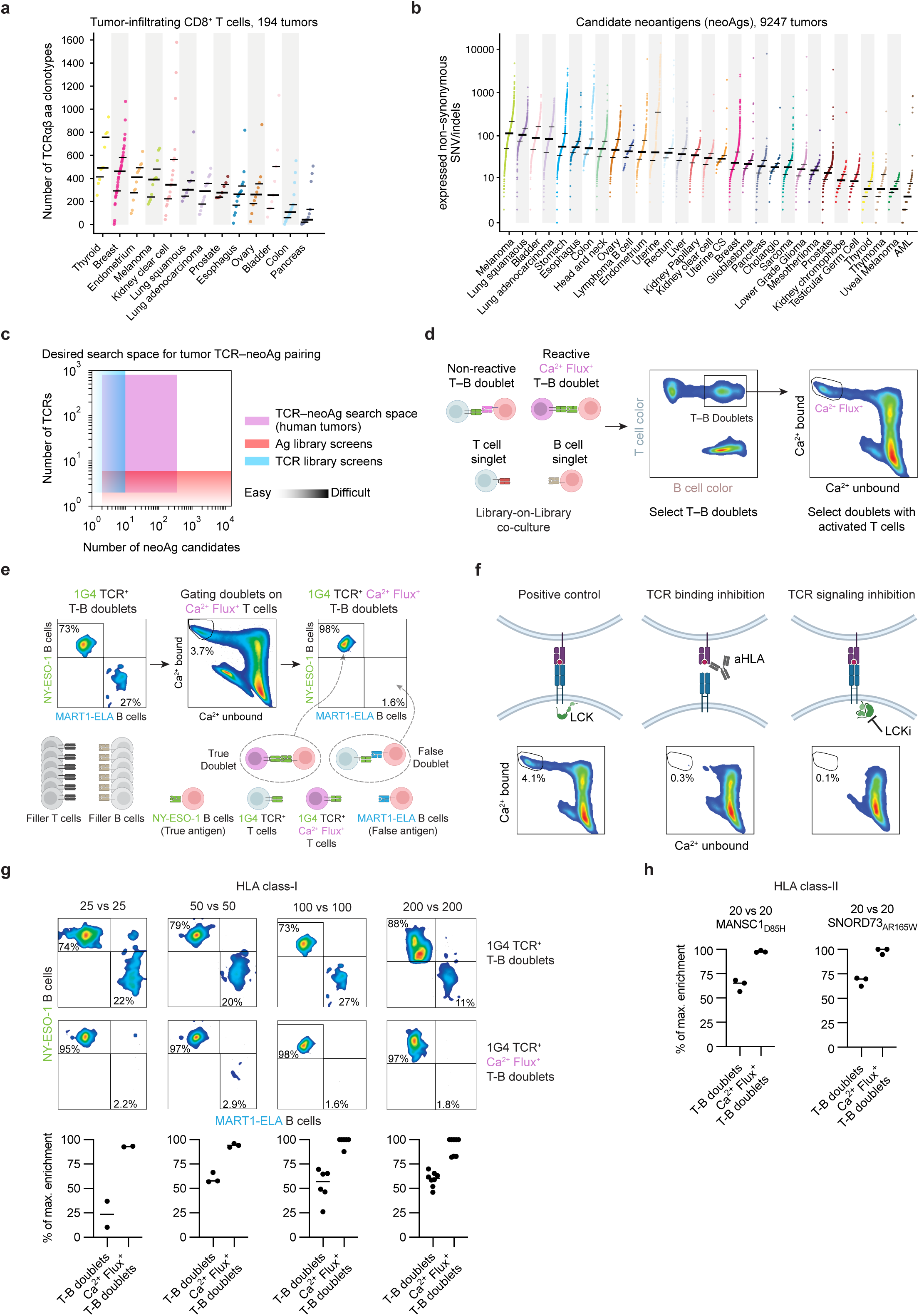
Interrogating the intratumoral TCR–neoantigen space using T cell–target cell doublets. **a)** Number of unique tumor–infiltrating CD8^+^ T cell TCRαβ amino acid clonotypes identified in 194 tumors by single-cell sequencing across 13 cancer types (studies listed in Supplementary Table 1). Dots represent individual tumors. Lines indicate the median and the 25th/75th percentiles of the number of clonotypes. **b)** Number of candidate neoantigens (neoAgs) across 31 cancer types in 9,247 TCGA tumor samples. Candidate neoAgs were defined as non-synonymous coding mutations (SNVs/indels) in expressed genes (>1 TPM). Mutations were detected by whole exome sequencing and gene expression by bulk RNA sequencing. Dots represent individual tumors. Lines indicate the median and the 25th/75th percentiles of the number of clonotypes. **c)** TCR–neoAg search space. Existing HLA-agnostic-^12,13,22^ and non-agnostic-^18–20,23^ cellular assays that can identify reactive TCR–neoAg pairs based on functional activity either screen reactivity of a single TCR against large antigen libraries^12,17–20^ (i.e., Ag library screens) or vice versa^13,21–23^ (i.e., TCR library screens), underscoring the need for HLA-agnostic functional library-on-library approaches that scale in both dimensions. **d)** Ca^2+^ flux^+^ T–B doublet isolation strategy. TCR library-transduced TCR-null Jurkat cells and neoAg-library-expressing autologous immortalized B cells are labeled with distinct CellTrace dyes. Jurkat cells are stained with indo-1, a fluorescent dye that shifts its emission spectrum when bound to Ca^2+^ (indo-1 Ca^2+^–bound^low→high^/indo-1 Ca^2+^–unbound^high→low^, Ca^2+^-flux^+^)^63^, to report proximal TCR signaling. Full gating strategy is depicted in Extended Data Fig. 2c. **e)** Evaluation of reactive T–B doublet formation in complex co-cultures. To simulate complex co-cultures, 1G4 TCR–transduced CD8^+^ TCR-null Jurkat cells and B cells expressing either cognate (NY-ESO-1) or control (MART1-ELA) antigen were admixed with parental filler cells at a defined ratio (e.g., 1:100). Labeling of B cells expressing cognate or control antigen with distinct fluorescent dyes was used to enable quantification of true-(1G4–NY-ESO-1) versus false (1G4–MART1-ELA) doublets. Selection of Ca^2+^-flux^+^ doublets enhances enrichment of reactive TCR–Ag pairs. Gating strategy illustrated in Extended Data Fig. 1g. **f)** Dependence of Ca^2+^ flux^+^ on TCR engagement. Schematic representation and representative flow cytometry plots (Gating strategy illustrated in Extended Data Fig. 1g) depicting the effect of disrupting TCR–peptide-HLA binding (anti-HLA antibody, aHLA) or proximal TCR signaling (Lck inhibitor; LCKi) on the proportion of T cell–B cell doublets displaying calcium flux at a 50×50 simulated screen complexity. **g)** Enrichment of reactive T–B doublets in complex co-cultures. Flow cytometry and enrichment quantification for HLA class I-restricted T cell–target cell pairs for the indicated combinatorial complexities. Gating strategy and calculation of percentage of maximum possible enrichment is illustrated in Extended Data Fig. 1g. Selection of Ca^2+^ flux^+^ T–B doublets increases signal purity across all tested complexities. **h)** Reactive doublet formation for HLA class II-restricted T cell responses. TCR-modified CD4^+^ TCR–null Jurkat cells and immortalized B cells expressing cognate or control antigen were admixed at a simulated combinatorial complexity of 20×20. Selection of Ca^2+^-flux^+^ doublets enhances enrichment of reactive TCR–antigen pairs.

Existing methods that can identify reactive TCR–neoAg pairs based on their functional activity in a single step either involve the screening of a single TCR against a library of candidate neoAgs^12,17–20^ (i.e., Ag library screens) or the reverse^13,21–23^ (i.e., TCR library screens) (Fig. 1c). While multiple such ‘one-on-library’ screens may be performed to increase the number of TCRs or neoAgs tested, the size of the TCR–neoAg search space (Fig. 1a-c) makes it necessary to develop conceptually distinct approaches.

To access the scale of the full human tumor TCR–neoAg space, we considered potential technologies in which reactivity of a library of candidate TCRs may be screened against a library of candidate neoAgs. To call reactive pairs in such library-on-library screens, the joint identification of TCR sequences and antigen sequences of cells that productively interact is critical. As TCR signaling induces enhanced adhesion between T cells and target cells^24,25^, we reasoned that isolation of T cell–target cell doublets from library-on-library co-cultures could form a means to identify functionally interacting pairs. Furthermore, as T cell activation induces a rapid increase in intracellular Ca^2+^ ^26^, we explored detection of cytosolic T cell Ca^2+^ levels as a second selectivity filter to increase the specificity of this approach. Importantly as sustained Ca²⁺ flux in conjugated T cells has been shown to correlate with TCR signal strength in T cell–target cell conjugates^27,28^, we reasoned that this approach could also provide a direct read-out of TCR potency (Fig.1d).

To evaluate whether this strategy allows the identification of reactive TCR–Ag pairs with high precision, we simulated a 100x100 complexity screen by diluting a reactive TCR–Ag pair and a non-reactive control TCR–Ag pair into ‘filler’ T and B cells. Specifically, T cells transduced with the HLA class-I-restricted NY-ESO-1 antigen-reactive 1G4 TCR were mixed with filler T cells, and B cells that expressed either the NY-ESO-1 antigen or a control (MART-1) antigen were mixed with filler B cells (Methods). To determine enrichment of true doublets that contained the reactive 1G4 TCR–NY-ESO-1 Ag pair over background, the relative abundance of 1G4 TCR–NY-ESO-1 and 1G4 TCR–MART-1 doublets was measured (Extended Data Fig. 1g). Quantification of 1G4 TCR T cell-containing doublets without concurrent analysis of Ca²⁺ signaling showed a reproducible but modest (∼3-fold) enrichment of true pairs over false pairs (73% versus 27%; Fig. 1e). Importantly, selection of 1G4 TCR^+^ T cell–B cell (T–B) doublets with active Ca²⁺ signaling resulted in 49-fold enrichment of true doublets over false doublets (98% vs 2% of total, Fig. 1e), indicating that selection of Ca^2+^ flux^+^ T–B doublets strongly enriches reactive TCR–Ag pairs over non-reactive pairs that are present at the same library input frequency. As controls of the specificity of this process, blocking of TCR binding to the NY-ESO-1-HLA-A*02:01 complex by anti-HLA antibody and inhibition of TCR signaling using an Lck inhibitor both abrogated Ca^2+^ signaling in cell doublets (Fig. 1f). Across simulated screening complexities (ranging from 625 to 40,000 simulated pairs), enrichment of true doublets in the Ca²⁺ flux^+^ T–B doublet gate remained consistently high, ranging from 93% to 100% of the maximally attainable cell pair enrichment (Fig. 1g and Extended Data Fig. 1g). In addition, co-culture of T cells modified with two different HLA class-II-restricted TCRs and their respective antigens demonstrated that this approach is also effective for the identification of HLA class II-restricted TCR–Ag pairs (Fig. 1h). Taken together, these results establish isolation of Ca²⁺ flux^+^ T–B doublets as a potential strategy to enable HLA allele-agnostic library-on-library screening.

### PAIR-Scan: HLA-agnostic library-on-library TCR–neoAg reactivity screens

Next, we explored assay conditions (See Methods and Extended Data Fig. 2) and data analysis strategies (see Methods and Extended Data Fig. 3) to develop **P**eptide **A**ntigen-**I**mmune **R**eceptor**-Scan** (PAIR-Scan), an HLA-agnostic library-on-library TCR–Ag functional screening platform (Fig. 2a). To evaluate the potential of PAIR-Scan to identify reactive TCR–antigen pairs in clinical samples, we focused on the intratumoral CD8^+^ TIL and candidate neoAgs from a metastatic melanoma patient Mel-Pt-A. In brief, we assembled a TCRαβ library of the 100 most expanded CD8^+^ TIL TCRαβ clonotypes of Mel-Pt-A using T-RAP^21^ high-throughput TCR synthesis, and introduced this TCR library into TCR-null CD8^+^ Jurkat reporter T cells^29^. In parallel, a minigene library of 100 Mel-Pt-A neoAg candidates with the highest tumor RNA expression was introduced into immortalized autologous B cells. To identify reactive TCR–neoAg pairs among the resulting 10,000 combinations, the Jurkat TCR library was exposed to the B cell neoAg library, individual Ca²⁺ flux⁺ T–B doublets were isolated (Extended Data Fig. 2b,c), and TCR and neoAg identity in individual cell doublets was determined by RNA-sequencing (Extended Data Fig. 3a–d). Expected background TCR–neoAg pairing frequencies, due to non-specific cell–cell interactions, were estimated by bulk sequencing of TCR and neoAg libraries (Extended Data Fig. 3a). PAIR-Scan confidence in the reactivity of each TCR–neoAg pair was then calculated as the probability of observing the frequency of that pair among isolated doublets, given its expected random pairing frequency (Extended Data Fig. 3e).

**Figure 2:**
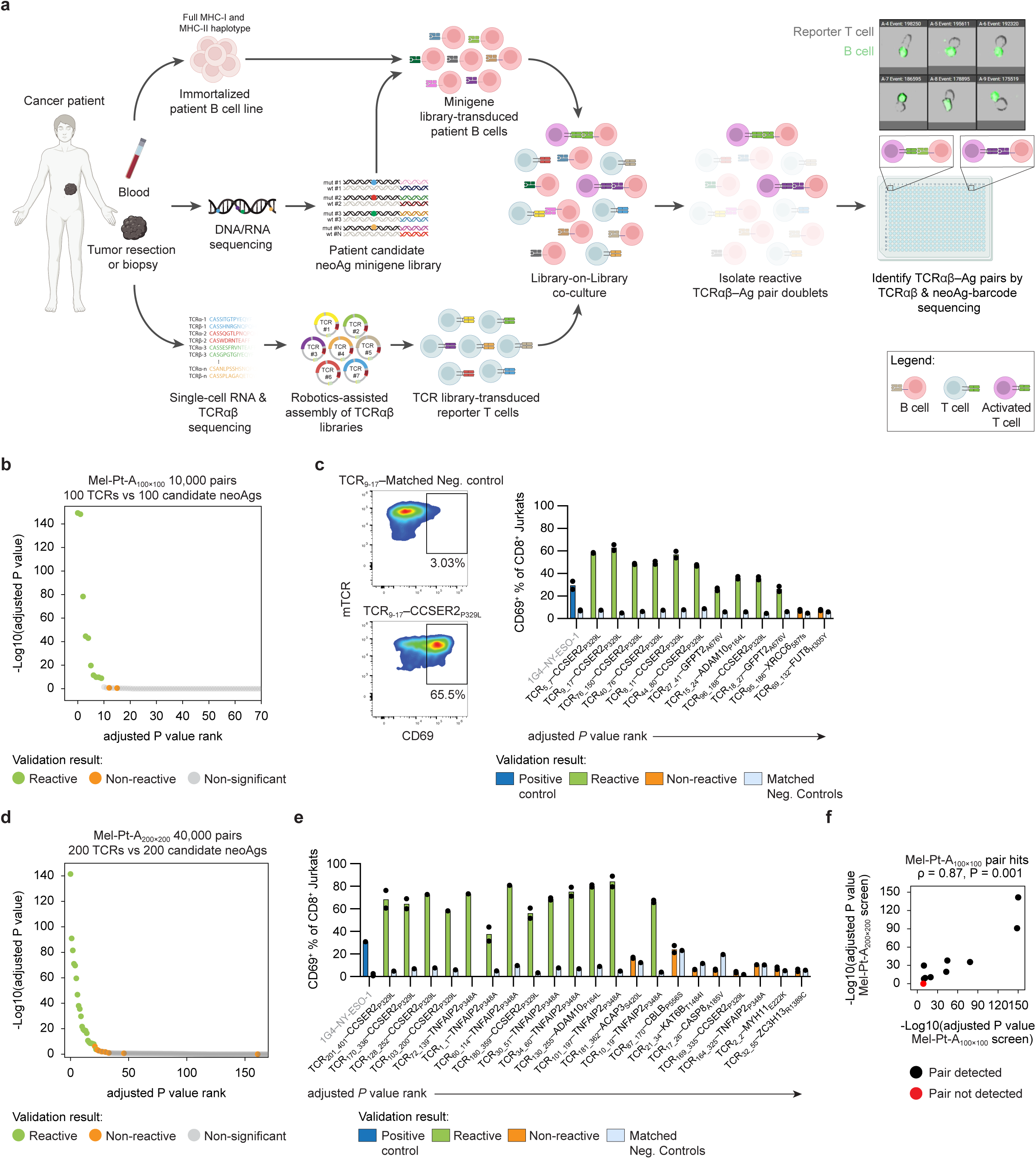
PAIR-Scan: Synthetic HLA-agnostic library-on-library TCR–Ag reactivity screening. **a)** Schematic of **P**eptide **A**ntigen–**I**mmune **R**eceptor**-Scan** (PAIR-Scan) TCR–Ag reactivity screening. DNA and RNA from tumor material is sequenced to identify candidate neoantigens (neoAgs) and their gene expression levels. Tumor-infiltrating lymphocyte (TIL) TCRs are identified by single-cell RNA and TCR sequencing and assembled as synthetic TCR libraries using T-RAP^21^. Patient-derived B cells, expressing the full range of patient HLA alleles, are immortalized by retroviral transformation with the anti-apoptotic factors Bcl-6/xL^64^ and transduced with personalized neoantigen (neoAg) libraries (Extended Data Fig. 2a). CD8^+^ or CD4^+^ TCR-null Jurkat cells are transduced with TCR libraries. The resulting T cell and B cell pools, expressing patient–matched TCRs and antigens, are co-cultured as described (Extended Data Fig. 2b), Ca^2+^ flux^+^ T–B doublets are sorted into 384-well plates (Extended Data Fig. 2c), and the identity of paired TCRs and neoAgs is determined by single-well sequencing (Extended Data Fig. 3a-d). PAIR-Scan confidence scores are assigned to observed TCR–neoAg pairs by comparing observed frequencies against expected random pairing frequencies, as estimated from original library distributions via bulk TCR and neoAg sequencing (Extended Data Fig. 3e, Methods). **b)** PAIR-Scan screen in melanoma patient A (10,000 combinations). PAIR-Scan confidence (- Log_10_ adjusted P values) in the reactivity of 10,000 screened TCR–neoAg combinations from melanoma patient A (Mel-Pt-A_100x100_). Dots, ranked by P value, represent putative TCR–neoAg pairs and are colored by functional validation status in (c). Green, reactive; orange, non-reactive; gray, non-significant. Note that P values are only shown for the first 70 out of 10,000 possible TCR–neoAg combinations **c)** Validation of TCR–neoAg pairs ranked highest by PAIR-Scan confidence in (b). Flow cytometry plots (left) and quantification of T cell activation (% CD69^+^ of CD8^+^ Jurkat cells, right) following individual co-cultures of indicated pairs from the Mel-Pt-A_100x100_ PAIR-Scan analysis. TCRs were tested against antigen-positive target cells (colored bars, *n* = 2 technical replicates) and matched negative control target cells (gray bars, *n* = 1 technical replicate). Positive control (1G4–NY-ESO-1) co-culture is depicted on the left. Full gating strategy is depicted in Extended Data Fig. 4a. **d)** PAIR-Scan screen in melanoma patient A (40,000 combinations). PAIR-Scan confidence in the reactivity of 40,000 screened TCR–neoAg combinations from melanoma patient A (Mel-Pt-A_200x200_). Dots, ranked by P value, represent TCR–neoAg pairs and are colored by functional validation status in (e). Green, reactive; orange, non-reactive; gray, non-significant. **e)** Validation of top TCR–neoAg pairs ranked by PAIR-Scan confidence in (d). Quantification of T cell activation (% CD69^+^ of CD8^+^ Jurkat cells) following individual co-cultures of newly identified pairs (i.e., not present in the Mel-Pt-A_100x100_ screen in (b,c)) with high PAIR-Scan confidence pairs in the Mel-Pt-A_200x200_ screen. TCRs were tested against antigen-positive target cells (colored bars, *n* = 2 technical replicates) and matched negative control target cells (gray bars, *n* = 1 technical replicate). Positive control (1G4–NY-ESO-1) co-culture is depicted on the left. Full gating strategy is depicted in Extended Data Fig. 4a. **f)** Reproducibility of TCR–neoAg identification across screens. Scatterplot comparing PAIR-Scan confidence in the reactivity of TCR–neoAg pairs with validating reactivity in Mel-Pt-A_100x100_ across Mel-Pt-A_100x100_ and Mel-Pt-A_200x200_ screens (two-sided Spearman’s ρ = 0.87, 95% CI: 0.41–1.00, P = 0.001). Black dots indicate pairs captured in both screens; red dot represents a single pair undetected in Mel-Pt-A_200x200_.

Analysis of the resulting data demonstrated highly significant enrichment for a number of TCR–antigen pairs. Importantly, the ten pairs ranked highest by PAIR-Scan confidence among the 10,000 screened TCR–neoAg pairs all showed validating reactivity in standard individual TCR–neoAg co-cultures, as assessed by activation-induced CD69 expression (Fig. 2b, c and Extended Data Fig. 4a). As a control, two lower-ranked pairs (13^th^ and 16^th^) both validated as non-reactive (Fig. 2b, c), indicating that PAIR-Scan confidence accurately separates reactive and non-reactive pairs. Seven of the ten reactive TCRs recognized the CCSER2_P329L_ neoAg, two were reactive to the GFPT2_A676V_ neoAg, and one TCR was identified for the ADAM10_P164L_ neoAg. To investigate whether PAIR-Scan can scale to higher complexity, we screened the reactivity of the 200 most expanded CD8^+^ TIL TCRs against the 200 most highly expressed neoAgs from the same patient (40,000 possible combinations, covering all genes with a TPM > ∼3). The top 21 TCR–neoAg pairs and the 23^rd^ pair by PAIR-Scan confidence in this screen all showed validating reactivity in arrayed testing (Fig. 2d,e). In contrast, the 22^nd^-ranked TCR–neoAg pair and all 7 pairs tested from the 24^th^-ranked pair onward all showed no reactivity in individual co-cultures, demonstrating a near perfect separation between reactive- and non-reactive TCR–antigen pairs (Fig. 2d,e). As compared to the 100×100 screen, five additional TCRs reactive to the CCSER2_P329L_ neoAg, one additional TCR reactive to the ADAM10_P164L_ neoAg, and seven TCRs reactive to the TNFAIP2_P348A_ neoAg were identified, underscoring that few neoAgs are recognized by a comparatively large number of different TCRs in this tumor. To investigate consistency of PAIR-Scan screens, we compared confidence values for the ten validated TCR–neoAg pairs identified in the 10K (100x100) screen with their corresponding confidence values in the 40K (200×200) screen. Confidence values for these ten pairs were highly correlated between independent screens (ρ = 0.87, P = 0.001), and nine out of ten pairs were again detected as hits in the 200×200 screen, with only the pair with the lowest confidence value in the 100×100 screen failing detection (Fig. 2f).

### PAIR-Scan identifies reactive TCR–neoAg pairs in high mutational load and T cell-infiltrated tumors

High mutational load tumors have been defined as those having approximately 150 or more nonsynonymous SNV/indels in expressed genes^1^. Given the relationship between ICB activity and tumor mutational burden^2,3^, analysis of tumor-specific T cell responses in high mutational load tumors is of particular interest. We therefore asked whether PAIR-Scan allowed the evaluation of all 781 neoAg candidates (approximately 98.2th percentile of TMB across tumors, Fig. 1b) identified in a metastatic lesion of a second melanoma patient Mel-Pt-B. To this purpose, we evaluated the reactivity of the 96 most clonally expanded CD8^+^ TIL TCRαβ clonotypes against all 781 neoAg candidates (74,976 combinations, Fig. 3a). PAIR-Scan confidence values were then used to select pairs for arrayed testing, using activation-induced CD137 expression on human peripheral blood CD8^+^ T cells as a readout. All five pairs ranked highest by PAIR-Scan confidence were confirmed to be reactive, while those ranked 7^th^, 9^th^, and 14^th^, included as controls, validated as non-reactive (Fig. 3b and Extended Data Fig. 4b). To also evaluate the feasibility of TCR-biased screens, we additionally evaluated the reactivity of all 393 CD8^+^ TIL TCRαβ clonotypes identified in a highly CD8^+^ T cell-infiltrated non-small cell lung cancer lesion of NSCLC-Pt-C against the 106 highest expressed neoAg candidates (41,658 combinations, Fig. 3c). In this screen, the top 4 TCRs ranked by PAIR-Scan confidence all showed validating reactivity against COL1A2_P771L_ neoAg, while the 5^th^ and 6^th^ ranked pair were confirmed to be non-reactive (Fig. 3d).

**Figure 3:**
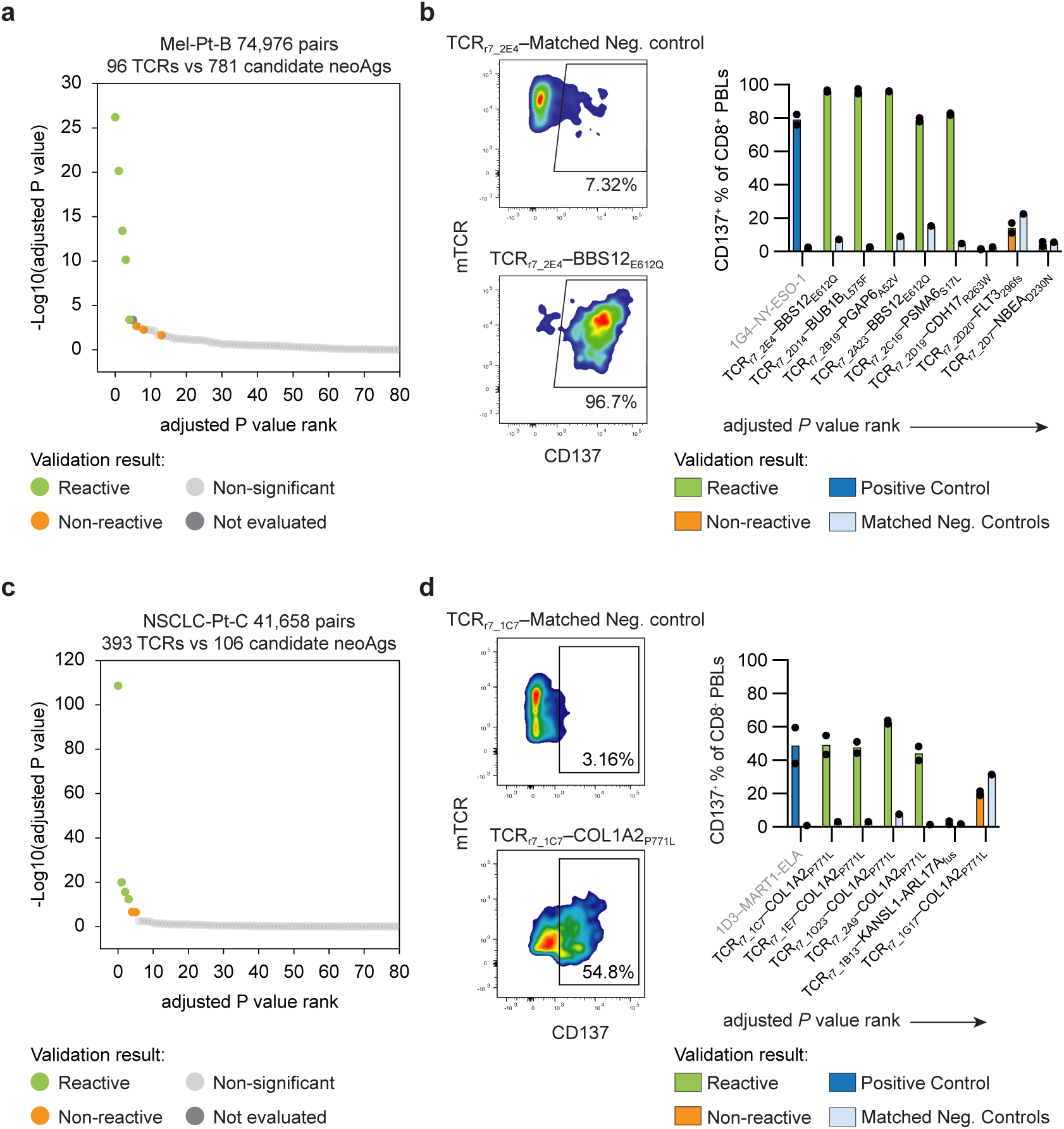
PAIR-Scan identifies reactive TCR–neoantigen pairs in high mutational load and T cell-infiltrated tumors. **a)** PAIR-Scan screen in melanoma patient B (74,976 combinations). PAIR-Scan confidence (–Log_10_ adjusted P values) in the reactivity of 74,976 screened TCR–neoantigen (neoAg) combinations from high mutational load tumor of melanoma patient B (96 TCRs vs. 781 candidate neoAgs; Mel-Pt-B). Dots represent TCR–neoAg pairs and are colored by functional validation status in (b). Green, reactive; orange, non-reactive; gray, non-significant. **b)** Validation of top TCR–neoAg pairs ranked by PAIR-Scan confidence in (a). Flow cytometry plots (left) and quantification of T cell activation (% CD137^+^ of CD8^+^ PBLs, right) in primary CD8^+^ T cells following individual co-cultures of candidate pairs from Mel-Pt-B. TCRs were tested against antigen-positive target cells (colored bars, *n* = 2 technical replicates) and matched negative control target cells (gray bars, *n* = 1 technical replicate). Positive control (1G4–NY-ESO-1) co-culture is depicted on the left. Full gating strategy is depicted in Extended Data Fig. 4b. **c)** PAIR–Scan screen in NSCLC patient C (41,658 combinations). PAIR-Scan confidence (–Log_10_ adjusted P values) in the reactivity of 41,658 screened TCR–neoantigen (neoAg) combinations from the highly CD8^+^ T cell-infiltrated tumor of NSCLC patient C (393 TCRs vs. 106 candidate neoAgs; NSCLC-Pt-C). Dots represent TCR–neoAg pairs and are colored by functional validation status in (d). Green, reactive; orange, non-reactive; gray, non-significant. **d)** Validation of top TCR–neoAg pairs ranked by PAIR-Scan confidence in (c). Flow cytometry plots (left) and quantification of T cell activation (% CD137^+^ of CD8^+^ PBLs, right) in primary CD8^+^ T cells following individual co-cultures of candidate pairs from NSCLC-Pt-C. TCRs were tested against antigen-positive target cells (colored bars, *n* = 2 technical replicates) and matched negative control target cells (gray bars, *n* = 1 technical replicate). Positive control (1D3–MART1-ELA) co-culture is depicted on the left. Full gating strategy is depicted in Extended Data Fig. 4b.

### PAIR-Scan identifies pairs of TCR and minimal peptides required for TCR reactivity modeling

For patient monitoring and for the design of e.g., personalized cancer vaccines, the definition of minigenes that encode T cell-recognized neoAgs is generally sufficient. However, for the training and evaluation of TCR reactivity prediction models, knowledge of the exact peptide sequence that is presented by HLA molecules is required^30^. We noted that the minigenes that encoded T cell-recognized neoAgs in patients A-C (Fig. 2 and 3) generally contained at least one predicted high affinity ligand for the patient HLA class I alleles (P=0.00001 relative to non-T cell-recognized neoAgs, Fig. 4a), suggesting that T cell reactivity could be screened directly against predicted minimal peptide binders. To explore this possibility, we asked whether a combination of computational peptide ranking and PAIR-Scan allowed identification of reactive TCR–minimal peptide pairs in an ultra-high mutational load tumor (Mel-Pt-D 3,879 mutations, approximately 99.6^th^ percentile of human tumors in Fig. 1b). Specifically, we ranked all 143,404 8-11-mer peptides containing these mutations by patient HLA allele binding confidence and mRNA expression (see Methods) and selected the top 500 ranked peptides for PAIR-Scan screening (50,000 combinations, Fig. 4b). PAIR-Scan analysis of this minimal peptide set against the 100 most expanded CD8^+^ TIL TCRαβ clonotypes from Mel-Pt-D revealed a number of hits. The four TCR–minimal peptide pairs ranked highest by PAIR-Scan confidence all validated as reactive, while the 5^th^, 6^th^, and 7^th^ ranked pairs all validated as non-reactive (Fig. 4b, c). Notably, within the set of 500 predicted HLA ligands that was analyzed experimentally, all the T cell-recognized minimal peptides ranked among the highest with respect to predicted HLA presentation (0.84 AUC, Fig. 4d,e), underscoring the value of rank-based selection of predicted minimal peptides. To evaluate whether rank-based selection may be used to enable PAIR-Scan analysis of TCR-minimal peptide pairs in larger patient groups, we re-evaluated a dataset of 70 patients from Rosenberg *et al.*^31^. In this dataset, selection of the top 500 ranked minimal peptides would suffice to include all experimentally validating T cell-recognized minimal peptides for a median of 100% of patients (mean 85%, Extended Data Fig. 5a). Together, these data suggest that PAIR-Scan analysis of predicted HLA ligands may be used to identify reactive TCR–minimal peptide pairs in human tumors with a low fraction of false negatives.

**Figure 4:**
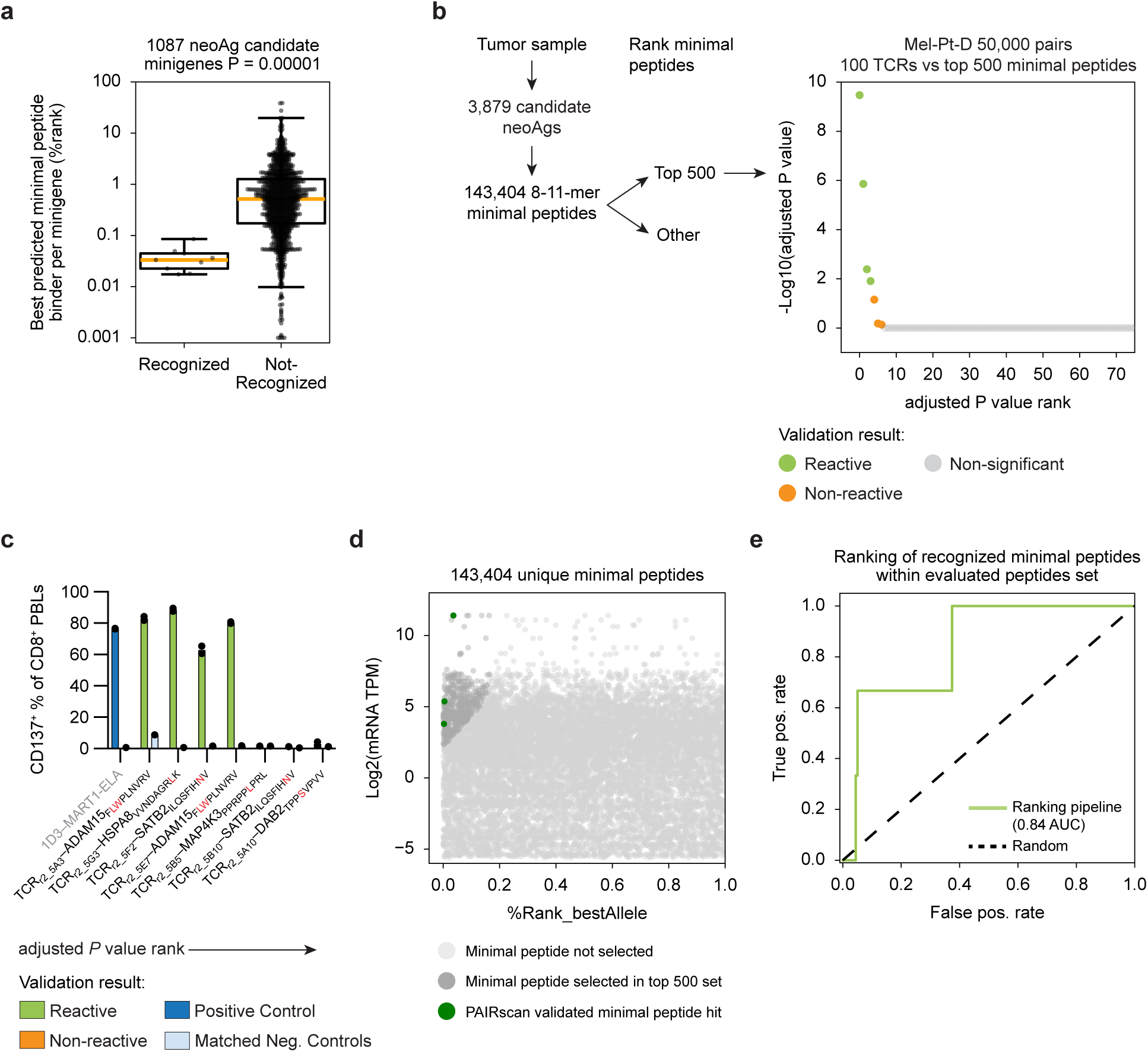
PAIR–Scan identifies TCR–peptide pairs required for TCR reactivity modeling. **a)** Predicted HLA class I binding scores for the best predicted minimal peptide binders derived from T cell-recognized and non-recognized neoantigen (neoAg) candidates in patients A-C PAIR-Scan screens. Comparison of the best–predicted binding %rank (MixMHCpred3.0) for patient-matched HLA class I alleles per neoAg-encoding minigene across screens (1,087 total minigenes, 9 T cell-recognized, 1,078 non-recognized). Peptides derived from T cell-recognized minigenes display significantly stronger predicted HLA binding (lower %rank) than peptides derived from non-recognized minigenes (two-sided Mann-Whitney U test, *U* = 615, *P* = 0.00001). **b)** Single-step identification of reactive TCR–minimal peptide pairs by PAIR-Scan screening. Selection strategy of minimal peptides (left) and statistical ranking (right) of PAIR-Scan confidence of 50,000 TCR–minimal peptide pairs from patient D (Mel-Pt-D). The top 500 minimal peptides based on predicted HLA binding confidence and mRNA expression, identified from a total of 143,404 candidates across 3,879 mutations, were screened against 100 most clonally expanded TCR amino acid clonotypes. Scatter plot displays PAIR-Scan confidence (–Log_10_ adjusted P values) in the reactivity of 50,000 screened TCR–peptide-MHC combinations. Dots represent TCR–minimal peptide pairs and are colored by functional validation status in (c). Green, reactive; orange, non-reactive; gray, non-significant. **c)** Validation of top TCR–minimal peptide pairs ranked by PAIR-Scan confidence in (b). Quantification of T cell activation (% CD137^+^ of CD8^+^ PBLs) in primary CD8^+^ T cells following individual co-cultures of indicated TCR–minimal peptide pairs from Mel-Pt-D. TCRs were tested against antigen-positive target cells (colored bars, *n* = 2 technical replicates) and matched negative control target cells (gray bars, *n* = 1 technical replicate). Positive control (1D3–MART1-ELA) co-culture is depicted on the left. Full gating strategy is depicted in Extended Data Fig. 4b. **d)** Candidate minimal peptide space of Mel-Pt-D. Predicted HLA class I binding rank (% rank of best allele, x–axis) and transcript abundance (log_2_TPM, y–axis) are depicted for all candidate 8-11-mer peptides with a < 1% predicted %rank. Light gray dots denote unselected minimal peptides, dark gray dots denote the top 500 candidates selected for screening, and green dots highlight the epitopes in PAIR–Scan validated hits. Note that PAIR-Scan validated hits are enriched for high predicted HLA binding even within the set of peptides that was selected for PAIR–Scan screening. **e)** Accuracy of prioritization of T cell-recognized minimal peptides (*n* = 3 unique recognized peptides) by HLA presentation prediction. ROC analysis demonstrating enrichment of functionally validated minimal peptides among the top–ranked candidates in Mel-Pt-D (AUC = 0.84, green line vs. random performance, dashed line).

Across the PAIR-Scan screens of patients A–D, all (13 out of 13) of the T cell-recognized neoAgs were derived from patient-specific mutations that were not recurrently observed in 9,247 tumor samples (TCGA), underscoring the need for technologies that can dissect neoantigen reactivity on a patient-specific basis. Furthermore, for 7 of these neoAgs, multiple reactive TCRs per neoAg were identified, thereby making it valuable to also understand the relative potency of these TCRs. Across screens, identified reactive TCR–neoAg pairs covered a range of low to high library frequencies (Extended Data Fig. 5b).

### PAIR-Scan identifies high potency TCR–neoAg pairs

The Mel-Pt-A 200×200 screen was performed twice to determine the reproducibility of PAIR-Scan. Both PAIR-Scan fold-enrichment and statistical confidence of the 12 CCSER2_P329L_ neoAg-reactive TCRs were highly correlated between these independent screens (fold-enrichment: ρ = 0.69, 95% CI: 0.05–0.99; significance: ρ = 0.83, 95% CI: 0.42–0.98; Fig. 5a). As prior work has shown that different TCRs that are reactive to the same (neo)Ag can vary substantially in their tumor killing potential^8–11^, we asked whether PAIR-Scan enrichment could predict TCR potency. To this purpose, we modified peripheral blood CD8^+^ T cells with seven different CCSER2_P329L_-reactive TCRs (Supplementary Table 2), covering a range of PAIR-Scan enrichment values, and compared their ability to kill CCSER2_P329L_-expressing B cells. Notably, the CCSER2_P329L_-reactive TCR set showed a wide variation in killing potential, as revealed by titration of available HLA class I molecules, and this killing potential was closely predicted by their PAIR-Scan screen enrichment values (ρ = 0.93, 95% CI: 0.41–1.0, Fig. 5b and Extended Data Fig. 6a). Furthermore, analysis of activation-induced IFNγ, TNFα, and cell surface CD107a expression for three of these TCRs with high, medium, and low PAIR-Scan enrichment values likewise showed a close match between PAIR-Scan enrichment and functional potency (Fig. 5c). Finally, differential killing activity that matched PAIR-Scan enrichment values was also observed when using CCSER2_P329L_-expressing D10 melanoma cells as target cells (Fig. 5d). To understand whether PAIR-Scan enrichment also correctly ranks TCRs reactive to other neoAgs, we evaluated two BBS12_E612Q_-reactive TCRs identified in Mel-Pt-B with medium-high (TCR_r7_2E4_) and medium-low (TCR_r7_2A23_) PAIR-Scan enrichment values, and three KIAA1671_A1129V_-reactive TCRs with high (TCR_r1_2B24_) and near-high (TCR_r1_2D24_ and TCR_r1_2G15_) PAIR-Scan enrichment values that were identified in a 5^th^ patient (Mel-Pt-E). For the two BBS12_E612Q_-reactive TCRs, a clear difference in killing activity was observed, and this difference was predicted by PAIR-Scan enrichment (Fig. 5e). For the 3 KIAA1671_A1129V_-reactive TCRs, strong killing activity was observed for each TCR, with relative potency still being predicted by PAIR-Scan enrichment (Fig. 5f).

**Figure 5:**
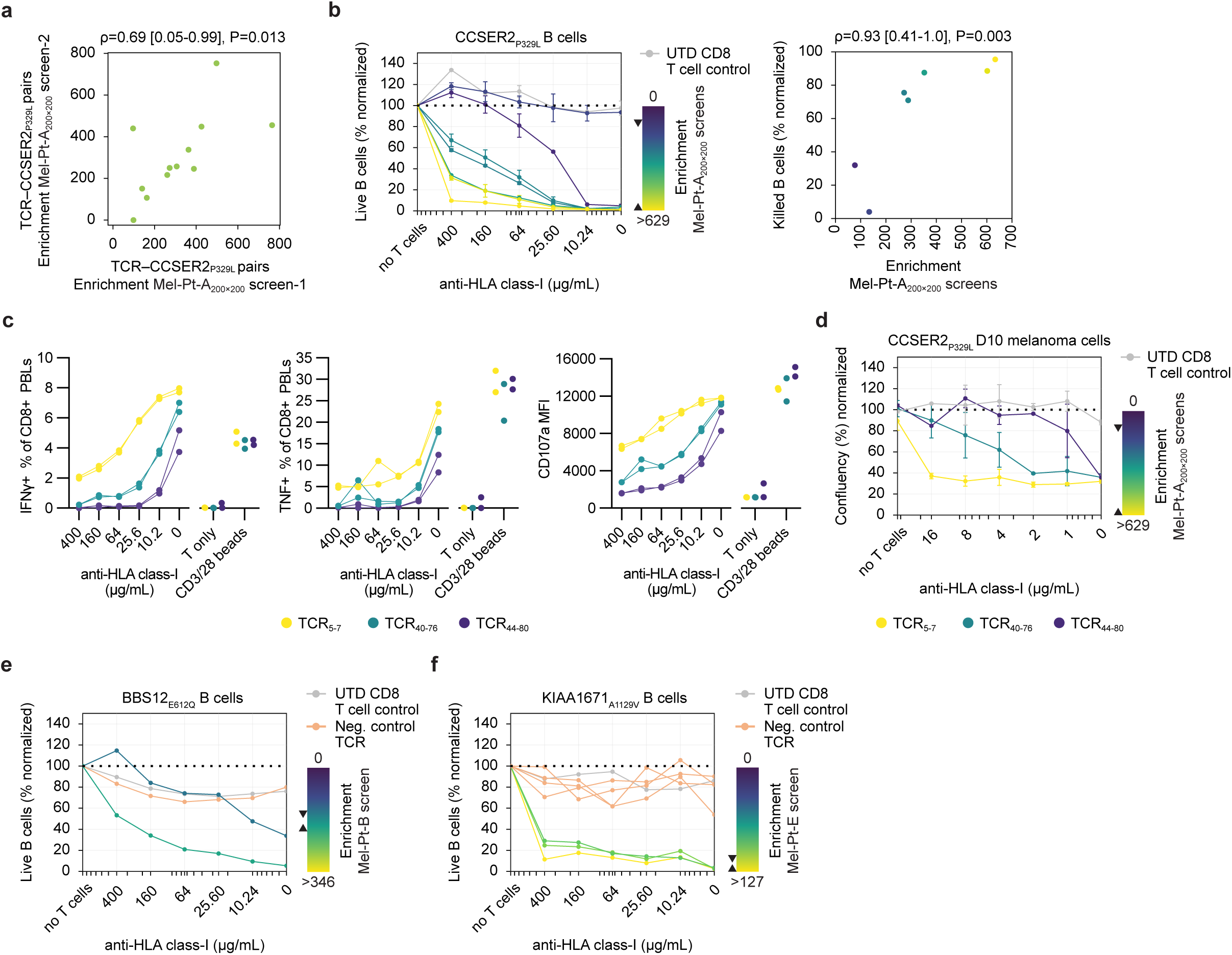
PAIR-Scan identifies high-potency TCR–neoAg pairs. **a)** Reproducible ranking of TCR–antigen pairs across independent PAIR-Scan screens. Fold-change enrichment of 12 CCSER2_P329L_-reactive TCRs in independent Mel-Pt-A_200x200_ PAIR–Scan screens (two-sided Spearman’s ρ = 0.69, 95% CI: 0.05–0.99, *P* = 0.013). Correlation of statistical significance (not depicted in figure) was ρ = 0.83, 95% CI: 0.42–0.98, *P* = 0.00079. **b)** Correlation between PAIR-Scan enrichment and T cell cytotoxicity. Cytotoxicity against CCSER2_P329L_-expressing B cells at the indicated anti-HLA class I concentrations is depicted for primary human T cells modified with seven CCSER2_P329L_-reactive TCRs (TCR_5–7_, TCR_9–17_, TCR_76–150_, TCR_8–11_, TCR_40–76_, TCR_44–80_, TCR_96–188_; ordered from highest to lowest killing efficiency) colored by their PAIR-Scan fold–change enrichment values (see Mel-Pt-A_200x200_ screens; lines show mean ± SD, *n* = 2 technical replicates). Normalized cytotoxicity closely correlates with PAIR–Scan screen enrichment values (right; two-sided Spearman’s ρ = 0.93, 95% CI: 0.41–1.0, *P* = 0.003, color scale = fold enrichment in PAIR-Scan Mel-Pt-A_200x200_ screens). Gating strategy and calculation of normalized cytotoxicity are depicted in Extended Data Fig. 6a. **c)** PAIR-Scan enrichment values rank TCRs by functional responsiveness. Degranulation (cell surface CD107a MFI) and intracellular cytokine staining (IFNg, TNFα) of primary human CD8^+^ T cells transduced with high-(TCR_5–7_), medium-(TCR_40–76_), or low-enrichment (TCR_44–80_) CCSER2_P329L_-reactive TCRs at the indicated anti–HLA class I concentrations (*n* = 2 technical replicates, shown as individual replicate lines). T cells only (T only) and anti-CD3/CD28 bead conditions serve as negative and positive controls. **d)** Correlation between PAIR–Scan enrichment and T cell cytotoxicity against CCSER2_P329L_-expressing melanoma cells. Imaging–based (Incucyte) quantification of CCSER2_P329L_-expressing D10 melanoma cell confluency following co-culture with primary human CD8^+^ T cells transduced with CCSER2_P329L_-reactive TCRs with high– (TCR_5–7_), medium-(TCR_40–76_), or low- (TCR_44–80_) PAIR-Scan enrichment scores at the indicated anti-HLA class I concentrations (lines show mean ± SD, n = 2 technical replicates). **e–f)** Correlation between PAIR–Scan enrichment and T cell cytotoxicity for BBS12_E612Q_- and KIAA1671_A1129V_-reactive TCRs. Cytotoxicity against BBS12_E612Q_- and KIAA1671_A1129V_-expressing target B cells at the indicated anti-HLA class I concentrations is depicted for primary human CD8^+^ T cells modified with BBS12_E612Q_- (TCR_r7_2E4_, medium high; TCR_r7_2A23_, medium low) and KIAA1671_A1129V_-reactive TCRs (TCR_r1_2B24_, high; TCR_r1_2D24_, near high; TCR_r1_2G15_, near high), colored by their PAIR-Scan screen enrichment values (*n* = 1 technical replicate). Negative control TCR (Neg. control TCR) and untransduced CD8^+^ (UTD CD8^+^ T cell control) control conditions are depicted. Gating strategy and calculation of normalized cytotoxicity are depicted in Extended Data Fig. 6a.

## Discussion

Here, we developed PAIR-Scan, a platform that identifies reactive TCR–Ag pairs among tens of thousands of candidate pairs and ranks these by their potency in a single screening step. PAIR-Scan is enabled by two distinct mechanisms: TCR-signaling dependent cell–cell adhesion that lowers the dissociation rate of reactive cell pairs, and enrichment of doublets with sustained T cell activation over transient non-specific doublets by simultaneous monitoring of intracellular Ca^2+^ flux.

While we exemplify the value of PAIR-Scan by profiling of T cell responses towards patient-specific cancer neoAgs, the approach that we describe may equally be used to identify TCR–Ag combinations for any other antigen that can be genetically encoded. For example, in related work we have performed PAIR-Scan screens for 9 melanoma, lung cancer, and bladder cancer patients, evaluating T cell reactivity against neoAgs, viral Ags, cancer germline Ags, and other tumor-associated Ags. In these screens, >50 confident reactive TCR–Ag pairs were identified, covering all the different Ag classes that were evaluated (data not shown). In view of the increasing evidence for a potential role of non-canonical antigens^32^, in future work it should also be of interest to use PAIR-Scan to screen for TCR reactivity against dark proteome-derived Ags, especially in tumors with a low mutational burden such as acute myeloid leukemia and pancreatic cancer. The capacity to rapidly identify reactive TCR–Ag pairs within a defined TCR and Ag space should likewise be valuable to support the ongoing development of personalized cancer vaccines^33^, e.g., by monitoring vaccine-induced TCR reactivity in peripheral blood. In addition to the screening of cancer- or vaccine-induced TCRs, PAIR-Scan may also be exploited to evaluate *in silico* designed TCRs or peptide-HLA-specific minibinders^34^. Beyond cancer, PAIR-Scan should be of value to dissect T cell reactivity in autoimmune disease. As a specific example, in type 1 diabetes, TCR clonotypes that bind self-peptide-HLA multimers but that show little functional activity against beta cells have been identified^35^ and large-scale functional screening of TCR–Ag pairs based on their potency would hence be of interest.

We demonstrate the ability of PAIR-Scan to identify reactive TCR–Ag pairs at a combinatorial complexity of up to 75,000 pairs and anticipate that PAIR-Scan can scale further, in particular with additional signal-to-noise optimizations. For example, a landing pad system^36^ that permits only a single TCR or Ag integration per cell will eliminate TCR-TCR and antigen-antigen co-occurrence in individual T or B cells and should thereby reduce noise. Ultimately, the scale of PAIR-Scan will be capped by the requirement for T cell – target cell encounter, an encounter probability that decreases in inverse proportion to the number of pairs screened. However, already at the current scale, PAIR-Scan can capture the TCR and neoAg space in a large fraction of human tumors. By focusing on cell–cell interactions, PAIR-Scan is distinct from RAPTR^37^ and ENTER-seq^38^, in which a peptide-HLA-expressing pool of lentiviral (LV) particles is incubated with TCR-expressing T cells. In addition, RAPTR and ENTER-seq require the separate production of peptide-HLA LV libraries for each of the 12 HLA class I and II alleles in each patient and, unlike for PAIR-Scan, it is unclear whether these approaches allow the direct scoring of TCR–peptide-HLA potency^37–39^.

It remains understudied to what extent tumor Ag-reactive TCRs differ in their potential to contribute to clinical tumor regression. In line with the fact that expression of inhibitory receptors is promoted by TCR signaling, potent antigen recognition can lead to more rapid T cell dysfunction^40^. At the same time, available clinical data provide greater evidence for insufficient, rather than excessive, potency as a primary limitation for tumor regression. In metastatic melanoma, patients who received T cells modified with a MART-1-reactive TCR with high *in vitro* potency showed a higher objective response rate than those who received a lower potency MART-1-reactive TCR that was obtained from the same TIL^8–10^. Consistent with this, an enhanced potency TCR reactive to the NY-ESO-1 cancer germline Ag has yielded high objective response rates in patients with metastatic synovial sarcoma and melanoma^41^. Further investigations into the effect of TCR–Ag potency on cancer regression in the human setting have been limited by the ability to identify TCRs with widely different activity towards the same tumor Ag. We anticipate that the capacity of PAIR-Scan to rank TCRs by their potency for the same tumor Ag will be valuable to address the relationship between receptor potency, T cell phenotype, and tumor regression. In addition, the set of seven TCRs with significantly different potencies towards the CCSER2_P329L_ neoAg will provide a useful model system for controlled studies of this effect (Supplementary Table 2). By making TCR–Ag pair potency measurable and rankable, PAIR-Scan may ultimately help guide the rational design of personalized immunotherapies, for instance by testing a mix of high- and intermediate-potency TCRs to balance short-term killing potential and longer-term engraftment in TCR-T therapy.

## Methods

### Patient material

Tumor tissue was collected from patients treated at NKI-AVL (Amsterdam, the Netherlands) with informed consent and in accordance with guidelines of the Medical Ethical Committee at the NKI-AVL. Tumor sample from Mel-Pt-A (referred to as "NKIRTIL063" in^12,13^) was processed as described^29^. For patients B-E, fresh tumor samples obtained by surgical resection were mechanically disrupted and digested for 4 h in RPMI 1640 medium (Thermo Fisher Scientific; #61870036) supplemented with 1 mg/mL collagenase type IV (Sigma-Aldrich; #C5138), penicillin-streptomycin (Roche; #11074440001), and 0.01 mg/mL pulmozyme (Roche). Resulting tumor digests were cryopreserved in liquid nitrogen.

### Antibodies

The following antibodies were used for flow cytometry: anti-CD3-APC (clone UCHT1, BD Biosciences #561811, 1:50 dilution); anti-CD4-FITC (clone RPA-T4, BD Biosciences #561005, 1:200 dilution), anti-CD4-FITC (clone OKT4, BioLegend #317408, 1:200 dilution), anti-CD4-TotalSeq-C (clone RPA-T4, BioLegend #300567, 1:5000 dilution); anti-CD8α-APC (clone SK1, BioLegend #344722, 1:50 dilution), anti-CD8α-AF700 (clone 3B5, Thermo Fisher Scientific #MHCD0829, 1:100 dilution), anti-CD8α-TotalSeq-C (clone SK1, BioLegend #344753, 1:500 dilution); anti-CD14-APC-H7 (clone MoP9, BD Biosciences #643077, 1:100 dilution); anti-CD16-APC-H7 (clone 3G8, BD Biosciences #560248, 1:100 dilution); anti-CD19-FITC (clone 4G7, BD Biosciences #340864, 1:200 dilution), anti-CD19-BV421 (clone HIB19, BioLegend #302234, 1:200 dilution), anti-CD19-BV711 (clone HIB19, BioLegend #302245, 1:100 dilution); anti-CD45-APC (clone HI30, BD Biosciences #555485, 1:50 dilution); anti-CD69-BV421 (clone FN50, BioLegend #310930, 1:50 dilution); anti-CD107a (clone H4A3, BioLegend #328619, 1:100 dilution); anti-CD137-BV421 (clone 4B4, BioLegend #309820, 1:50 dilution); PE-conjugated anti-mouse TCRβ constant domain (clone H57-597, BD Biosciences #553172, 1:150 dilution); PerCP-Cy5.5-conjugated anti-human TCRαβ constant domain (clone IP26, BioLegend #306724, 1:100 dilution); anti-TNFα-PE (clone Mab11, BioLegend #502908, 1:150 dilution); anti-IFNγ-FITC (clone B27, BioLegend #506504, 1:100 dilution); mIgG1-TotalSeq-C (clone MOPC-21, BioLegend #400187, 1:250 dilution), anti-human Hashtag-TotalSeq-C (clone LNH-94, BioLegend #394661, 1:100 dilution). Live/Dead Fixable Near-IR Dead Cell Stain (Thermo Fisher Scientific), Propidium Iodide (PI), or DAPI was used to identify live cells. Data from flow cytometry experiments were acquired using FACSDiva (v 8.0.2, BD Biosciences) or FACSChorus (v 6.1, BD Biosciences) software and analyzed using FlowJo (v 10.7.1, BD Biosciences).

### Isolation of tumor-infiltrating T cells and patient-derived B cells

Tumor-infiltrating T cells were isolated from tumor digest, and patient-derived B cells were isolated either from tumor digest or peripheral blood mononuclear cells (PBMCs). Patient PBMCs were isolated from peripheral blood by Ficoll-Paque (GE Healthcare; #17-1440-03) density gradient. Cryopreserved tumor digests were thawed at 37 °C in 4 °C RPMI 1640 medium supplemented with 10% human serum (Sigma-Aldrich; #H3667), penicillin-streptomycin, and Benzonase (Millipore; #70746; 1:1000 dilution), and washed with PBS containing 0.04% BSA at 4 °C. For patient A, tumor-infiltrating CD8^+^ T cells (DAPI^-^CD45^+^ CD3^+^ CD8^+^) were sorted from cryopreserved tumor digest and B cells (IR-Dye^−^ CD3^−^ CD14^−^ CD16^−^ CD19^+^) were sorted from patient PBMCs, as described in^12,13^. For patients B–E, each tumor sample was split into three equally sized aliquots, and each aliquot was labeled with a unique hashtag-TotalSeq-C antibody, enabling identification of T cell doublets co-encapsulated within the same gel bead-in-emulsion (GEM) (see Single-cell RNA & TCRαβ sequencing). Aliquots were blocked with Human TruStain FcX (BioLegend; #422301; 1:20 dilution) and stained in MACS buffer with anti-CD3-APC, anti-CD4-FITC, anti-CD8-AF700, anti-CD19-BV421, anti-CD4-TotalSeq-C, anti-CD8α-TotalSeq-C, mIgG1-TotalSeq-C, and a unique hashtag-TotalSeq-C antibody per aliquot. Propidium iodide viability dye was added immediately prior to sorting. Tumor-infiltrating CD8^+^ T cells (PI^-^CD3^+^ CD8^+^ CD4^-^) and B cells were sorted (PI^-^ CD3^-^ CD19^+^) using a FACSAria Fusion cell sorter (BD Biosciences).

### Cell lines

Isolated patient-derived B cells were stimulated for 24-36 h with 50 ng/mL human recombinant IL-21 (BioLegend; #571206) and irradiated (55 Gy) hCD40L^+^ murine L cells in IMDM (Thermo Fisher Scientific; #12440061) containing Penicillin-streptomycin and 10% heat-inactivated fetal bovine serum (Thermo Fisher Scientific; #10270-106). Subsequently, B cells were transduced with retrovirus encoding human Bcl-6, Bcl-xL, and a GFP or NGFR transduction marker using Retronectin (Takara; #T100A), as per the manufacturer’s instructions. Cells were re-stimulated weekly with 50 ng/mL IL21 and bi-weekly with fresh irradiated L cells.

TCRα^-^β^-^Jurkat cells (ATCC; #TIB-152), expressing either human CD8αβ or CD4-zeocinR, as previously described^29^, were cultured in RPMI 1640 medium supplemented with penicillin-streptomycin and 10% heat-inactivated fetal bovine serum.

A Jurkat reporter cell line expressing a fusion protein composed of human CD3z (Uniprot P20963) and enhanced green fluorescent protein (eGFP) was generated by means of retroviral transduction. The CD3z and eGFP encoding sequences were linked by an oligonucleotide sequence encoding a G_4_SG_4_S-linker, ordered as a gene fragment (Twist Bioscience) and cloned into pBabe_hygro (Addgene #1765) using restriction digest.

Human primary CD8^+^ T cells were obtained from healthy donor-derived PBMCs by Ficoll-Paque density gradient and magnetic bead separation (Miltenyi Biotec; #130-096-495) and cultured in RPMI 1640 medium supplemented with penicillin-streptomycin, 10% heat-inactivated human serum (Sigma-Aldrich; # H3667, and 30 U/mL human IL-2 (Proleukin).

D10 melanoma cells (CVCL_H945; gift from the Daniel Peeper lab) were cultured in DMEM (Thermo Fisher Scientific; #11965092) supplemented with penicillin-streptomycin and 10% heat-inactivated fetal bovine serum.

### Single-cell RNA G TCRαβ sequencing

Single-cell RNA & TCRαβ sequencing of patient A (“NKIRTIL063”) tumor material is described in^13^. For patient B-E, sorted tumor-infiltrating CD8^+^ T cells were washed twice in PBS with 0.04% BSA at 4 °C, live T cells were counted, and 33 µL of 1200 T cells/µL (∼39,600 cells) were loaded into a Chromium single-cell sorting system (10x Genomics) for single-cell RNA and TCRαβ sequencing using the Chromium Next GEM Single Cell 5’ and V(D)J Reagent Kit v2 (10X Genomics) and sequenced on Illumina NovaSeq6000 system.

### Exome and RNA sequencing

Whole-exome sequencing (WES) and RNA sequencing (RNA-seq) were performed as previously described^12,29^. Briefly, genomic DNA and RNA were co-extracted from formalin-fixed, paraffin-embedded tumor material using the AllPrep DNA/RNA FFPE kit (Qiagen; #954734). Genomic DNA from patient PBMCs was purified using the DNeasy Blood & Tissue kit (Qiagen; #69504). Exomes were enriched using the SureSelect XT2 Human All Exon V6 kit (Agilent), and strand-specific RNA-seq libraries were prepared with the TruSeq Stranded mRNA sample preparation kit (Illumina), both following the manufacturers’ protocols. Libraries were sequenced on Illumina HiSeq 2500 or NovaSeq 6000 platforms.

### TCR library cloning

TCR libraries were assembled using the T-RAP high-throughput TCR assembly platform^21^. T-RAP assembles TCRs in 384 well plates, allowing the generation of highly uniform custom TCR libraries by pooling, while simultaneously creating an arrayed TCR archive from which individual TCR hits can be picked for validation. In brief, to demultiplex CDR3-J-encoding oligonucleotide pools (Twist Bioscience) into CDR3α-Jα- and CDR3β-Jβ fragments of defined TCRs in individual wells, well-specific orthogonal (non-interacting) primers were used to obtain double-stranded (dsDNA) using PCR amplification. Vectors encoding all human TRAV, TRBV, and mouse TCR constant regions were described previously. CDR3-J, TRAV, TRBV, and TCR constant sequences were combined into the viral T-RAP vector (Extended Data Fig. 2a) using Golden Gate assembly. Following Golden Gate assembly, reaction products were pooled to generate TCR libraries and purified, as previously described^21^. Purified Golden Gate pools were transformed at >1,000x coverage into electrocompetent Endura E.coli (Biosearch Technologies; #60242-2) by electroporation (10 uF, 600 W, 1800 V) with a Bio-Rad Pulser Xcell, using 75 ng of DNA per 25 µL of electrocompetent cells. Plasmid DNA was isolated by means of column purification (Qiagen; #12143).

### Candidate neoAg and minimal peptide library cloning

Candidate neoAg and minimal peptides were cloned into the P20 (HLA class I-restricted epitope screens) or P12 (HLA class II-restricted epitope screens) lentiviral expression vectors (will be deposited at Addgene). Candidate neoAg and minimal peptide-encoding library dsDNA fragments were generated by PCR using 0.66 ng library oligonucleotide pool (Twist Bioscience), 0.02 U/µL Q5 polymerase (NEB; #M0493L), 0.25 µM of both forward and reverse plate-specific primers, 0.2 mM dNTPs (Sigma-Aldrich; #11969064001), and 1x Q5 polymerase buffer in a total volume of 100 µL, followed by incubation in a thermocycler (30 s at 98 °C, 11 cycles of 10 s at 98 °C, 20 s at 65 °C and 35 s at 72 °C, followed by 2 min at 72 °C). PCR product was purified using Monarch PCR & DNA Cleanup Kit (NEB; #T1130S) and PCR product yield was determined using an Agilent Bioanalyzer 2100 (DNA 7500 kit). Golden Gate assembly was performed using 300 ng P20 or P12 lentiviral plasmid DNA and neoAg-encoding or minimal peptide-encoding library fragments at a 1:3 molar ratio of vector to insert. Reactions contained 100 U/µL T4 ligase (NEB; #M0202M), 0.5 U/µL BsmBI-v2 (NEB; #R0739L), and 1× T4 ligase buffer in a total volume of 80 µL, and were incubated in thermocyclers for 35 cycles of 10 min at 42 °C and 10 min at 16 °C, followed by 10 min at 55 °C and 15 min at 75 °C for heat inactivation. Golden Gate products were purified, electroporated at >1,000× library coverage, and used for plasmid DNA purification as described for TCR libraries.

### Viral transductions

Retrovirus was produced by transfecting FLY-RD18 packaging cells with retroviral transfer plasmid DNA encoding individual TCRs or TCR libraries (T-RAP pMX-TCRαβ-puroR; Extended Data Fig. 2a) using Lipofectamine 3000 Reagent (Thermo Fisher Scientific). Retrovirus encoding the eGFP-CD3zeta fusion protein (pBabe_eGFP-CD3zeta_hygroR) was produced similarly. Lentivirus was generated by transfection of 293T cells (Lenti-X cells, Takara; #632180) with lentiviral transfer plasmid DNA encoding individual neoAgs, minimal peptides or neoAg libraries (P20- or P12-puroR; Extended Data Fig. 2a), together with helper plasmids encoding pGALV-MTR envelope (Addgene; #163612)^42^ and lentiviral packaging proteins (Addgene; #12260). Production of retrovirus used for B cell immortalization was previously described^12^.

Suspension cells (Jurkat cells, primary T cells, immortalized B cells, primary B cells) were transduced by means of spinfection using retronectin. Primary T cells were stimulated with anti-CD3/28-coated beads (LifeTechnologies; #11131D) and 150 U/mL IL-2 (Proleukin) for 48 h prior to transduction. B cells were stimulated for 24-36 h with 50 ng/mL human recombinant IL-21 and irradiated (55 Gy) hCD40L^+^ murine L cells prior to transduction. For spinfection, non-treated 6-well (Jurkat TCR or neoAg B cell libraries) or 24-well plates were coated with retronectin and blocked with 2% BSA (Sigma-Aldrich; #A4737) according to the manufacturer’s instructions. Subsequently, 0.5-1 mL of filtered viral supernatant was added and centrifuged for 90 min at 2000 g. Next, viral supernatant was removed and 1.5M cells (6-well, Jurkat TCR or neoAg B cell libraries transductions) or 0.25-0.5M cells (24-well transductions) were added per well. After 24-72 h, cells were harvested and transduction efficiency was assessed by flow cytometry using an anti-mouse TCRβ constant domain antibody for T cells or mKelly1 fluorescence as a marker of P20 or P12 transduction for B cells. Transductions with TCR- and neoantigen libraries were performed at a transduction rate of 2.5–5% to minimize the occurrence of multiple integrations and at a library coverage of >1000x.

Adherent cells (D10 melanoma cells) were transduced by addition of filtered viral supernatant containing 4 mg/mL polybrene (Sigma-Aldrich; #TR-1003-50UL) for 24 h, after which viral supernatant was replaced with fresh culture medium.

48–72 h post-transduction, cells underwent selection with antibiotics. Jurkat cells and primary T cells transduced with T-RAP pMX-TCRαβ-puroR were selected in 2.5 µg/mL puromycin (Thermo Fisher Scientific; #A1113803) for 48 h. Jurkat cells transduced with pBabe_eGFP-CD3zeta_hygroR were selected in 500 µg/mL hygromycin B (Invivogen; #ant-hg-1) for 10 days. Immortalized B cells and D10 melanoma cells transduced with P20 or P12 were selected in 2.5 µg/mL puromycin for 48 h. Transduction and selection efficiency were monitored by flow cytometry. In case library transduction efficiency was below 75% at 48–72 h after the first puromycin selection for Jurkat cells or at 6 days after the first puromycin selection for B cells, a second selection round with 5 µg/mL puromycin was performed for 48 h.

### Experimental modeling of screening parameters

To identify optimal conditions for detection of reactive TCR–Ag pairs using doublet formation, CD3z:eGFP^+^ CD8α^+^β^+^ TCR-transgenic Jurkat-null cells (i.e., 1G4-TCR^+^ Jurkat cells) were co-cultured with target-matched (i.e., NY-ESO-1^+^) and unmatched (i.e., MART1-ELA^+^ or GLC^+^) immortalized B cells, labeled with CellTrace Far Red and Yellow (Thermo Fisher Scientific; #C34572, #C34567; 1:20,000 dilution) respectively, at a 1:1 ratio. To simulate library diversity, TCR-transgenic Jurkat cells and antigen^+^ target cells were each mixed with parental Jurkat cells (“filler T cells”) and parental B cells (“filler B cells”), at the indicated ratios prior to co-culture initiation. In addition, all B cells were stained with anti-CD19-BV711, filler T cells were stained with anti-human TCR-PerCP-Cy5.5, and all T cells were stained with Indo-1 AM dye (200 ng/mL, Invitrogen; #I1203) for 30 min at 37 °C. To initiate co-cultures, stained cells (1M total B cells and T cells) were mixed at the indicated ratios in 5 mL polystyrene flow cytometry tubes, centrifuged twice (5 min, 200 g, room temperature), and then incubated (5–15 min, 37 °C) without dissociation of cell pellets (Extended Data Fig. 2b). Directly prior to centrifugation steps, as well as flow cytometry, tubes were briefly flicked and passed through a 35 µm filter to dissociate cell pellets.

The effect of assay parameter variation on the enrichment of true doublets over false doublets was determined by measuring the number of true and false T cell–B cell doublets after gating on T cell–B cell conjugates (CD19^+^TCRb^+^ *OR* CD19^+^eGFP^+^) that showed calcium flux (indo-1 Ca^2+^-bound^hi^, indo-1 Ca^2+^-unbound low) (gating and enrichment calculation are shown in Extended Data Fig. 1g). Flow cytometric analysis was performed on a BD S8 FACS discovery or Sony ID7000. True doublet enrichment was further assessed in the presence of Lck-inhibitor (8 µM, Sigma-Aldrich; #428205) and pan-MHC-class I blocking antibody (160 µg/mL; Biolegend #311427).

### PAIR-scan screening

To reduce cell density-associated activation, Jurkat TCR libraries were cultured at a concentration below 500,000 cells per mL for a period of 7 days prior to screening. Four days before screening, the fraction of CD8ab^+^ muTCRβ^+^ Jurkat T cells and P20 transduced B cells was monitored by flow cytometry using an anti-mouse TCRβ constant domain antibody for T cells and mKelly1 fluorescence as a marker of P20 or P12 transduction for B cells. Three days before screening, puromycin was removed from Jurkat TCR library culture medium, as Ca^2+^ flux can be reduced after puromycin treatment^43^.

At the day of PAIR-Scan screening, Jurkat TCR libraries were stained with fluorescent dye (CellTrace Yellow, 1:20,000 dilution) and Indo-1 AM dye (200 ng/mL) for 30 min at 37 °C. The GFP^-^ B cell libraries of Mel-Pt-A and NSCLC-Pt-C were additionally stained with fluorescent dye (CellTrace Far Red, 1:20,000 dilution) for 30 min at 37 °C. The GFP^+^ B cell libraries of Mel-Pt-B, D, and E were not stained with CellTrace.

To start co-cultures, 625,000 Jurkat TCR library and neoAg library B cells were passed through a 35 µm filter and mixed in a 5 mL polystyrene flow cytometry tube (Extended Data Fig. 2b). Next, the tube was centrifuged for 5 min at 200 g and 37 °C to promote cell–cell contact, and cells were incubated for 5-15 min at 37 °C in pellet form to allow TCR–signaling-induced adhesion. Cells were subsequently redistributed by flicking the tube to dissociate transient non-specific contacts while retaining stable reactive T–B cell doublets, and centrifugation and incubation steps were repeated once. After the second round of incubation, the tube was flicked, cells were passed through a 35 µm cell strainer and immediately sorted into 384-well plates containing one-step RT-PCR buffer (see single-well sequencing), using the cell gating strategy described in Extended Data Fig. 2c. Dead cell exclusion with this gating was based primarily on forward/side scatter morphology. As an additional step, SYTOX Red Dead Cell Stain (Thermo Fisher Scientific; #S34859, 1:100 dilution) can be added prior to the first round of centrifugation, with removal of unbound dye by replacement with warm medium. Given that doublets eventually dissociate and that both Ca^2+^ flux and cell – cell adhesion function optimally at 37 °C, the sample chamber of the cytometer was set to 37 °C and every 32 min a fresh co-culture was started. The sorter collection stage (plate) was set to 4-8 °C. Directly after cell sorting, plates were centrifuged at 2000 g for 2 min at 4 °C and directly incubated in a thermocycler for single-well sequencing.

### Plate-based single-well paired TCR-and Ag-barcode-sequencing

Individual T–B cell doublets were sorted into 384-well plates (DNA library preparation strategy illustrated in Extended Data Fig. 3a) containing 0.48 µL one-step reverse transcription (RT)-PCR mix, 0.8 µL of 6 µM barcoded RT primers (unique barcode per well; Supplementary Table 3), and a 3 µL overlay of filtered silicone oil (Sigma-Aldrich; #317667-1L) to prevent evaporation. The one-step RT-PCR mix consisted of 0.5 µM tcr_Fw, 0.5 µM pep_Fw, and 0.5 µM R1_Rev primers (Supplementary Table 4), 0.1% Triton X-100 (Sigma-Aldrich; #T8787-50ML), and 1X LunaScript Enzyme in LunaScript buffer (NEB; #E1555L). Plates were prepared by dispensing the well-barcoded RT primers using a Microlab STAR 384 liquid handler (Hamilton), immediately followed by the silicone oil overlay. RT-PCR mix was then dispensed into each well using an I.DOT dispenser (Dispendix, BICO). Plates were centrifuged at 1200 g for 1 min at 4 °C and stored at -80 °C up to six months before use.

For PAIR-Scan screens, 384-well plates were thawed on ice and centrifuged at 1200 g for 1 min at 4 °C immediately prior to sorting. After sorting of T–B cell doublets, plates were sealed (Bio-Rad; #MSF1001), centrifuged at 2000 g for 2 min at 4 °C, and incubated in a C1000 384-well thermocycler (Bio-Rad; #1851138). RT-PCR products were generated by RT for 15 min at 55 °C followed by PCR, using 1 min at 98 °C, 25 cycles of 10 s at 98 °C, 30 s at 65 °C and 75 s at 72 °C, followed by 2 min at 72 °C. Following RT-PCR, plates were centrifuged at 1000 g for 1 min at room temperature. PCR products of each plate were pooled into pooling reservoirs containing 1 mL pre-added silicone oil (Clickbio; #VBLOK200) by centrifugation at 250 g for 1 min, and the aqueous phase of PCR reaction products was manually recovered from the reservoir. Pooled PCR products were purified using 1.1X AMPure XP beads (Beckman Coulter; #A63881), washed three times with 80% ethanol, and eluted in 30-35 µL nuclease-free water. Concentration and quality of purified plate PCR products were assessed using an Agilent Bioanalyzer 2100 (DNA 7500 kit).

PCR products of individual plates were dual-indexed using an Illumina sample index PCR by combining 10.5 ng plate PCR product, 0.5 µM p5 and p7 sample index primer (Supplementary Table 5), 0.2 mM dNTPs, 0.02 U/µL Phusion HF enzyme in 1X Phusion HF buffer (Thermo Fisher Scientific; #F530L) in a total volume of 100 µL, followed by PCR (3 min at 98 °C, 7-10 cycles of 10 s at 98 °C, 30 s at 55 °C and 1 min at 72 °C, followed by 5 min at 72 °C). Sampled index PCR products were purified using 0.9X AMPure XP beads, washed three times with 80% ethanol, and eluted in 15-25 µL. Concentration and quality of purified plate PCR products were assessed using an Agilent Bioanalyzer 2100 (DNA 7500 kit). Sample-indexed PCR product of each plate was pooled at equimolar ratio and sequenced using an Illumina MiSEQ (single read, 150 bp) at a coverage of at least 2000 reads per well (768K reads per plate).

### Bulk TCR and Ag-barcode library sequencing and library QC

Bulk TCR and Ag-barcode library sequencing of library plasmid DNA and library cell lines (DNA library preparation strategy shown in Extended Data Fig. 3a) was used for library quality control (QC) and to estimate the abundance of individual TCRs and Ags. This abundance estimate was used to calculating the expected frequency of TCR–Ag pairs under random pairing in PAIR-Scan screen analysis (see below). QC assessed the percentage of successfully assembled TCRs and Ags, the percentage of sequence-perfect TCRβ molecules (i.e., exact matches with no mismatches or indels), and the uniformity of the TCR and Ag libraries. T cell and B cell libraries were lysed at >1,000x library coverage in DirectPCR Lysis Reagent (Viagen) containing 500 µg/mL proteinase K in a total volume of 62.40 µL by incubation at 55 °C for 60 min, 85 °C for 30 min, and 94 °C for 5 min, and lysis product was diluted by adding 40 µL nuclease-free water. TCRβ chains were dual-indexed by Illumina sample index PCR using 0.5 µM TCR_p7 primer (Supplementary Table 5) annealing to the T-RAP retroviral vector (pol region), 0.5 µM TCR_p5 primer annealing to the murine TCRβ constant domain (Supplementary Table 5), 1x NEBNext HiFi 2x PCR mastermix (NEB; #M0541L), 1 ng purified plasmid library DNA or one or more aliquots of 12 µL diluted lysis product in a total volume of 30 µL. Reactions were incubated in a thermocycler for 30 sec at 98 °C, followed by 14 cycles for plasmid library DNA or 22–25 cycles for lysis product, of 10 s at 98 °C, 20 s at 72 °C and 50 s at 72 °C, and 2 min at 72 °C. PCR products were purified using 0.7x AMPure XP beads (Beckman Coulter), washed three times with 80% ethanol, and eluted in 20 µL nuclease-free water, and product concentrations were measured using an Agilent Bioanalyzer 2100 (DNA 7500 kit).

Ag DNA barcodes were dual-indexed by Illumina sample index PCR using 0.5 µM P20_p7 or P12_p7 primer annealing to the lentiviral vector P20 5′ UTRs or P12 promoter (Supplementary Table 5) and 0.5 µM P20_P12_p5 primer annealing to a truncated non-coding sequence of the murine TCRβ constant domain (Supplementary Table 5) inserted downstream of the DNA barcode to create a shared RT handle with the T-RAP vector (Extended Data Fig. 2a). PCR reactions contained 1× NEBNext HiFi 2× PCR Master Mix and 10.5 ng purified plasmid library DNA in a total volume of 30 µL. PCR was performed in a thermocycler (30 s at 98 °C, followed by 11 cycles of 10 s at 98 °C, 20 s at 72 °C, and 50 s at 72 °C, followed by 2 min at 72 °C). PCR products were purified using 0.75× AMPure XP beads (Beckman Coulter), washed three times with 80% ethanol, and eluted in 35 µL nuclease-free water. Product concentrations were measured using an Agilent Bioanalyzer 2100 (DNA 7500 kit).

All library cell lines were QC’ed on the day of screening. For Jurkat TCR library lines, the median percentage of library TCRs that was detected was 98.6%, and the median fold difference in TCR abundance between the 5th and 95th percentiles was ∼14-fold (Supplementary Table 6). For B cell neoAg library lines, the median percentage of library neoAg-barcodes that was detected was 100%, and the median fold difference in neoAg abundance between the 5th and 95th percentiles was ∼10-fold (Supplementary Table 6)

### TCR–Ag pair validation and cytotoxicity assays

Reactivity of T cells transduced with individual TCRs was analyzed by flow cytometry after a 6-8 h (Jurkats) or 12-16 h (human peripheral blood CD8^+^ T cells) co-culture with target cells expressing the indicated Ag. Co-cultures were performed in U-bottom 96-well plates at an effector-to-target ratio of 1:1. T cells incubated in the absence of target cells with or without phorbol 12-myristate 13-acetate (50 ng/mL; Sigma-Aldrich) and ionomycin (1 µg/mL; Sigma-Aldrich) addition served as positive and negative controls, respectively. Cells were stained with IR-Dye and anti-mouse TCRβ constant domain antibody, in combination with anti-CD8 and anti-CD137 (when using primary T cells) or anti-CD69 (when using Jurkat cells), and subsequently analyzed by flow cytometry. Co-culture with B cells expressing irrelevant epitopes (matched negative control B cells in the figures) was used to determine background T cell activation levels. For experiments in which intracellular cytokine production was measured, protein transport inhibitor cocktail (eBioscience; #00-4980-03) was added 3 h after co-cultures were started. Cells were then fixed and permeabilized following the manufacturer’s instructions (Biolegend; #426803), and intracellular TNFα and IFNγ were measured by flow cytometry using anti-TNFα and anti-IFNγ. To measure T cell degranulation, anti-CD107a was added at the start of co-cultures.

The ability of peripheral blood CD8^+^ T cells transduced with individual TCRs to kill target cells presenting different levels of available Ag was analyzed in 48 h co-cultures. To determine TCR potency, the number of available HLA class I molecules was titrated using an anti-human HLA-A,B,C antibody (Biolegend; #311441). Untransduced T cells served as negative controls. Where indicated, T cells transduced with irrelevant TCRs were included as additional negative controls. B cell killing assays were performed in U-bottom 96-well plates at an effector-to-target ratio of 2:1 to 4:1. Target B cells expressing Ag were mixed with untransduced parental B cells colored with Cell Trace Yellow that served as an internal negative control at a 1:1 ratio. Cells were stained with DAPI viability dye just before analysis by flow cytometry. B cell cultures without added T cells served as negative controls. The gating strategy and calculation of T cell cytotoxicity is depicted in Extended Data Fig. 6a. D10 melanoma killing assays were performed by live-cell imaging (IncuCyte S3, Sartorius) in flat-bottom 96-well plates at an effector-to-target ratio of 1:1. D10 cells were seeded at 10,000 cells per well. The next day, CD8^+^ T cells were added, and confluency was analyzed over 48 h using IncuCyte S3 Software (v2018B). D10 cells cultures without added T cells served as negative controls.

### Single-cell RNA G TCRαβ sequencing analysis and TCRαβ library design

Cell Ranger (patient A v3.0.0, patient B and & v8.0.1, patient D and E v7.1.0) was used to process Chromium Single Cell 5′ V(D)J sequencing data to generate single-cell RNA expression profiles and to assemble and annotate TCRs. For patient D and E, a custom human IMGT-based VDJ reference was used, and for patient A, B and C, the 10X Genomics-provided refdata-cellranger-vdj-GRCh38-alts-ensembl-7.1.0 was used. Cells with a high proportion of mitochondrial transcripts (patient B and C >7.5%, patient D and E >25%) or low total unique molecular identifier (UMI) counts (patient B and C < 600 UMI counts, no UMI threshold applied for patient D and E because EmptyDrops was run in Cell Ranger) were excluded. For patient B–E, cells with high isotype control antibody-derived tag (ADT) counts were first removed, sample demultiplexing and doublet removal based on hashtags was performed using HTODemux (Seurat^44^; patient A v4.2.1, patient B and C v5.1.0, patient D and E v4.3.0), and CD8^+^ and CD4^+^ T cells were annotated by gating on a scatterplot of CD8 versus CD4 ADT counts.

To call and count TCRαβ amino acid clonotypes, the filtered_contig_annotations.csv generated by CellRanger were parsed using the TCRtoolbox python package. Briefly, contigs were filtered on Cell Ranger’s high confidence, is_cell, full-length, and productive flags. Barcodes with only a single chain, or with multiple chains that were all alpha or all beta but not both, were removed. For patient B and C, dual TCRα and dual TCRβ chains were filtered per cell barcode, separately for the alpha and beta chains. Filtering was based on the ratio of the non-top chain’s UMI count to the top chain’s UMI count. Dual TCR chains that passed this filter were then ordered deterministically by UMI count when building the clonotype string, to prevent incorrectly counting dual chain clonotypes twice. For each 10X dataset, a non-stringent ratio threshold was set by inspecting the density-normalized histogram of non-top chain UMI count ratios, choosing a cutoff just below the point where non-top alpha chains are less frequent than non-top beta chains (for patient B and C, a ratio cutoff of 0.3), as dual alpha chains are generated more frequently than dual beta chains^45^. Amino acid clonotypes were defined by TRAV, TRAJ, TRBV, and TRBJ gene usage and the CDR3α and CDR3β amino acid sequences. Amino acid clonotype counts and frequencies were calculated separately for CD8^+^ and CD4^+^ T cells of each sample by counting of cell barcodes. To create TCRαβ clonotypes for screening from cells expressing dual alpha and/ or dual beta chains, cell barcodes were expanded into multiple rows that covered the two (one dual chain) or four (two dual chains) possible combinations, with the counts of the parent dual chain clonotype being used for the newly generated single pair clonotypes.

For TCRαβ library design, the resulting unique amino acid clonotypes were sorted by their clonotype frequency and the indicated top clonotypes were selected for screening and prepared for T-RAP^21^ TCR assembly using TCRtoolbox software.

### Number of candidate TCRαβ clonotypes in single-cell RNA G TCR-sequencing studies

The single-cell RNA & TCR-sequencing datasets used for analysis are described in Supplementary Table 1. To ensure consistency across datasets, raw sequencing data had to be available for processing with Cell Ranger, or raw Cell Ranger outputs generated using a version >3.0.0 had to be available. Second, datasets either had to contain existing T-cell lineage annotations (i.e., CD4⁺ or CD8⁺) or raw sequencing data that could be used to generate such annotations. Where lineage annotations were available, these were used directly. For datasets without existing annotations, raw data were processed using Cell Ranger 9.0.1 using default parameters, including cell calling with EmptyDrops. Cells with a high proportion of mitochondrial transcripts (>15% of total UMI counts) or fewer than 100 detected genes (non-stringent as EmptyDrops already filtered low quality cells) were also excluded. For datasets with available CD8 and CD4 ADT counts, CD8^+^ and CD4^+^ T cells were annotated in the same way as in “Single-cell RNA & TCRαβ sequencing analysis and TCRαβ library design”. For datasets with only gene expression data, T cells were annotated using a T cell score (*CD3D, CD3E, TRAC, TRBC1 and TRBC2*) computed with the Scanpy score_genes function, and non T cells were removed. CD8^+^ T cells were then annotated by setting a threshold based on a *CD8A* and *CD8B* score histogram. As *CD4* is poorly detected in single-cell RNA sequencing datasets, CD4^+^ T cells were annotated as non-CD8^+^ T cells. Cells that had both a high *CD8A* and *CD8B* score and high CD4 expression were removed.

The filtered_contig_annotations.csv from all datasets were parsed using the TCRtoolbox as described in “Single-cell RNA & TCRαβ sequencing analysis and TCRαβ library design”, except that dual TCR chains were filtered using a fixed threshold of 0.4 for the ratio of the non-top chain’s UMI count to the top chain’s UMI count for all datasets. To balance the number of tumors analyzed per cancer type, the breast cancer dataset from Nederlof et al. (Supplementary Table 1) was randomly downsampled to match the size of the second-largest cancer dataset.

### Number of neoAg candidates in TCGA tumor samples

Tumor tissue gene expression data for TCGA were retrieved from the TCGA data portal as transcripts per million (TPM), quantified against the GENCODE v36 annotation^46^. Somatic mutations for the same cohorts from whole-exome sequencing against matched normal samples (GDC MuTect2 workflow, variant-aggregated and masked) were downloaded from the UCSC Xena Browser^47^. Expression and mutation tables were linked by the 12-character TCGA patient barcode. In cases where a donor contributed multiple RNA aliquots, the first was retained.

For each donor we defined the expressed tumor mutational burden as the number of non-synonymous coding somatic variants occurring in a gene that was transcriptionally active in that tumor. Somatic calls were restricted to those passing all filters (FILTER == PASS) and to protein-altering consequences, defined as any of missense_variant, frameshift_variant, stop_lost, start_lost, inframe_insertion, inframe_deletion, protein_altering_variant, coding_sequence_variant, splice_acceptor_variant, or splice_donor_variant. HGNC gene symbols in the mutation tables were mapped to Ensembl gene identifiers using the GENCODE v36 GTF (one identifier per symbol), and variants without a mapping were discarded. Identical variant calls (donor, chromosome, position, reference and alternate allele) were collapsed to one event per donor.

Each retained variant was paired with the TPM of its gene in the matching donor’s tumor RNA, and was scored as expressed if TPM > 1. Only donors with paired mutation and expression data were analyzed, yielding 9,247 donors across 31 cancer types.

### Exome and RNA sequencing data analysis

Exome and RNA sequencing data were analysed as previously described^12,29^. Exome sequencing reads were aligned to the GRCh38 human reference genome using BWA^48^ (v0.7.10). For patient A, germline variants were called with GATK HaplotypeCaller^49^ and somatic variants with GATK MuTect2^50^ (GATK v3.7.0), annotated with SnpSift^51^ (v4.3p). For patients B-E, the same alignment and variant-calling steps were performed using the nf-core/sarek pipeline^52^ (v3.1.2; BWA 0.7.17 and GATK v4.3.0), with annotation by SnpSift^51^ (v5.1). For patient A, RNA sequencing reads were aligned with STAR^53^ and mRNA expression was quantified with Salmon^54^. For patients B-E, RNA sequencing reads were similarly aligned with STAR and quantified with Salmon using the nf-core/rnaseq pipeline^52^ (STAR v2.7.10a, Salmon v1.10.1); RNA gene fusions were additionally called using Arriba^55^ (v2.3.0) via the nf-core/rnafusion pipeline^52^.

### Design of candidate neoAg and minimal peptide libraries

Tumor-specific variants included as a source of candidate neoAgs consisted of expressed non-synonymous SNVs, in-frame insertions (up to 20 inserted amino acids), in-frame deletions (up to 30 deleted amino acids), and frameshifting indels and SNVs that resulted in loss of stop codons. Frameshifts or stop-loss variants with up to 40 novel downstream amino acids to the new stop codon were included for patients A-C. All frameshifts and stop-loss variants were included for patients D and E. Variants for patients A, C, and D were selected by mRNA expression TPM rank, while all variants with TPM > 0 were screened for patient B and E.

SNV-encoding minigenes were designed as sequences encoding the mutated amino acid flanked by 15 amino acids upstream and downstream (31-mers). In-frame insertions were encoded as the inserted amino acids flanked by 14 amino acids upstream and 15 amino acids downstream (31–48-mers). Novel junctions created by in-frame deletions were flanked by 15 amino acids upstream and 15 amino acids downstream (30-mers, patients B-E) or 16 and 15 amino acids (31-mers, patient A). For patients A-C, frameshifting indels and stop-loss SNVs were encoded as minimally 15 amino acids of wild-type sequence (extended further upstream when the novel downstream sequence left the minigene shorter than 31 amino acids) followed by up to 40 amino acids of novel downstream sequence to the new stop codon (31–55-mers). For patient E, wild-type upstream and novel downstream sequences were segmented into 31-mers with 15-amino-acid overlap without a length limit on the novel downstream sequence (see below). For patient D, minimal peptides were required to overlap with the novel downstream sequence for library inclusion. RNA fusion transcripts were translated *in silico* and encoded as 15 amino acids upstream and 15 amino acids downstream of the junction between the two fused gene products. Minimal peptides were defined as all 8-11-mer peptides containing at least one mutated residue, for SNVs, in-frame insertions, and frameshifting indels. For in-frame deletions, minimal peptides were required to contain both amino acids forming the neojunction created by the deletion. Binding confidence (%Rank) to all patient HLA class I alleles was predicted using MixMHCpred3.0^56^. Peptides with %Rank_bestAllele >2% were filtered to reduce the minimal peptide sorting set size. Protein TPM mRNA expression and %Rank_bestAllele were converted to percentile ranks, where 1.0 represents the highest TPM and lowest %Rank_bestAllele. A combined sort score was calculated as 0.2 × TPM percentile rank + 0.8 × %Rank_bestAllele percentile rank, and minimal peptides were sorted by this score. Duplicate minimal peptide amino acid sequences were then removed, retaining the first-ranked sequence. The top 500 minimal peptides, as based on weighted predicted MHC class I binding and RNA expression, were selected for PAIR-Scan screening.

NeoAg-encoding minigene and minimal peptide-encoding library oligonucleotide pools were prepared with the python package TCRtoolbox. NeoAg and minimal peptide amino acid sequences were codon optimized with the TCRtoolbox codon optimizer. Briefly, codon optimization removed type IIS BsmBI recognition sites, four-base Golden Gate overhangs, Kozak sequences, primer binding sites, repeats longer than four nucleotides, and terminator motifs (TTTTT or AAAAA). In addition, codon optimization enforced a global GC content between 35–60%, as previously described^21^. A TAA stop-codon was directly added after each codon-optimized coding sequence and followed by a unique 18 bp (neoAg minigene libraries) or 36 bp (minimal peptide libraries) DNA barcode from^57^. The resulting nucleotide sequences were flanked by BsmbI recognition sites and orthogonal primer binding sites (stored as constants in the TCRtoolbox software) for PCR amplification. To identify design errors before wet lab cloning, cloning of each neoAg or minimal peptide was modeled *in silico* using pydna^58^, modeling both restriction digestion, DNA ligation, and PCR.

### PAIR-Scan screen data analysis

To provide an end-to-end computational pipeline in which screen preparation and screen analysis share the same underlying information, the TCRtoolbox software that performs both screen preparation and PAIR-Scan screen sequencing data analysis was used.

#### Counting TCR and Ag-barcode transcript UMIs per well

TCR and Ag-barcode transcript UMIs were counted per well using the TCRtoolbox count_umi_cell function. Well DNA barcodes and UMIs were extracted with umi_tools^59^ (v 1.1.6), and reads were trimmed with cutadapt^60^ (v 5.1), removing 3’ bases below Phred 31 together with adapters and constant TCR and vector sequences, such that only the diverse, high-quality sequence remained for alignment. Using TCRtoolbox as an end-to-end pipeline, the adapter and constant-region trim sequences, along with the alignment reference, can be read directly from the files created during screen preparation or from constants stored in the software. Trimmed reads were aligned to a combined TCR and Ag-barcode reference with BWA MEM^61^ (v 0.7.19). Alignments were sorted and indexed with samtools (v 1.22) and filtered with pysam (v 0.23.3), retaining alignments with 0 mismatches, 0 indels, and a minimum overlap of 102-107 bp for TCR and 36 bp for Ag-barcode references. Retained alignments were deduplicated by UMI and counted per well and per transcript using umi_tools count, yielding a well-by-transcript UMI count table.

#### Counting bulk TCR and Ag-barcode reads

Bulk TCR and Ag-barcode reads were counted per well with the TCRtoolbox count_reads_bulk function. Bulk TCR and Ag-barcode libraries were trimmed, aligned, and filtered with the same cutadapt, BWA MEM, samtools, and pysam based steps described above (0 mismatches, 0 indels, minimum overlap 106 bp for TCR and 36 bp for Ag-barcode), except that aligned reads per reference were counted directly with pysam, yielding each TCR’s and Ag-barcode’s read abundance. Ag-barcode read counts were then combined into a single count per Ag by summing counts for the same Ag.

#### Counting detected TCR–Ag pairs

Detected pairs per well were called from the UMI count table in three steps, as shown in Extended Data Fig. 3c. TCR and Ag-barcode transcripts were first filtered on a minimum UMI count. If a well contained multiple TCR or multiple Ag-barcode transcripts, non-top transcripts were filtered separately within each transcript type based on their ratio to the well-specific top transcript, using a threshold that was depth-scaled on the top transcript UMI count. Every remaining pairwise TCR–Ag combination per well was then called as a detected pair, and the number of wells for each pair was counted.

#### Epi-Epi and TCR-TCR co-occurrence filtering

Epi-Epi and TCR-TCR co-occurrence filtering (algorithm shown in Extended Data Fig. 3d) was used to remove TCR–Ag pairs with insufficient independent well support for their TCR, Ag, or both.

#### Calculating PAIR-Scan confidence and TCR–Ag pair enrichment values

Expected random TCR–Ag pairing frequencies were estimated from bulk TCR and Ag-barcode library sequencing, as illustrated in Extended Data Fig. 3e. Bulk TCR and Ag read counts were each normalized to sum to 1 and the expected frequency of each TCR–Ag pair due to random pairing was calculated as the product of these values. The observed frequency of each pair was calculated as its well count divided by the total number of wells with at least one detected TCR–Ag pair. Screen fold-enrichment was calculated as the observed frequency divided by the expected frequency due to random pairing. PAIR-Scan confidence in a pair’s reactivity was the probability, under a one-sided binomial test, of observing at least its well count *k* among the total number of sorted wells with at least a detected TCR–Ag pair *n*, given its expected pairing frequency *p*, implemented with Scipy^62^ (scipy.stats.binom.sf; v 1.15.2). Resulting *P* values were corrected for multiple testing with Benjamini-Hochberg FDR correction, adapted from the statsmodels package. Only the most significant pair for TCRs that showed high PAIR-Scan confidence with more than one unique Ag was reported.

## Data availability

All PAIR-Scan screen sequencing data will be deposited in the National Center for Biotechnology Information’s Sequence Read Archive.

## Code availability

The TCRtoolbox python package is available at: https://github.com/schumacherlab/TCRtoolbox, including a TCR library parsing and design example notebook: https://github.com/schumacherlab/TCRtoolbox/blob/main/notebooks/tcr_parsing_examples/parse_and_design_tcr_library_example.ipynb, a TCR library assembly preparation tutorial: https://github.com/schumacherlab/TCRtoolbox/blob/main/tutorials/assembly_tutorial.md, an Ag library design example notebook: https://github.com/schumacherlab/TCRtoolbox/blob/main/notebooks/epi_assembly_examples/design_ag_library_example.ipynb, a minimal peptide library design example notebook: https://github.com/schumacherlab/TCRtoolbox/blob/main/notebooks/epi_assembly_examples/design_minimal_peptide_library_example.ipynb, an Illumina TCRβ chain or TCRα chain bulk-sequencing read counting tutorial: https://github.com/schumacherlab/TCRtoolbox/blob/main/tutorials/count_bulk_illumina_seq_reads_cli_tutorial.md, a PAIR-Scan single-well paired TCR-and neoAg-barcode-sequencing counting tutorial: https://github.com/schumacherlab/TCRtoolbox/blob/main/tutorials/pair_scan_counting_tutorial.md, and a PAIR-Scan analysis example notebook: https://github.com/schumacherlab/TCRtoolbox/blob/main/notebooks/sequencing_analysis/pair_scan_analysis_example.ipynb. Code used to generate figures in this paper is available at: https://github.com/schumacherlab/schumacherlab_publications_code/tree/main/messemaker_and_richterich_et_al_2026.

## Acknowledgements

This work was supported by the Louis Jeantet prize and Stevin Award (to T.N.S.). This work was also delivered as part of the MATCHMAKERS team, supported by the Cancer Grand Challenges partnership funded by Cancer Research UK (CGCATF-2023/100003), the National Cancer Institute (OT2CA297206) and the Mark Foundation for Cancer Research, and as part of the ILLUMINE team, supported by the Cancer Grand Challenges partnership funded by Cancer Research UK (CGCATF-2025/100002), the National Cancer Institute, the Cancer Research Institute and KiKa (Children Cancer Free Foundation). J.H. was supported by EMBO (ALTF 643-2025) and Marie Skłodowska-Curie Actions (101208803). Research at the Netherlands Cancer Institute is supported by institutional grants from the Dutch Cancer Society and the Dutch Ministry of Health, Welfare and Sport. We would like to thank Michael Dustin (University of Oxford, UK), Peter Friedl (RadboudUMC, The Netherlands), and members of the Schumacher lab for advice and scientific input.

## Author information

T.N.S., C.A.R., M.Me., and W.S. conceived and designed the study. J.B.A.G.H. provided patient material. J.H. and S.d.P. quantified the number candidate TCR and neoAgs in tumor samples across cancer types. Ž.M., W.S., and R.V. performed tumor exome, RNA, and TIL single-cell RNA & TCR sequencing experiments. M.Me., C.A.R., J.G., J.U., B.M., Ž.M., A.G., and M.Mu. assembled TCR and Ag libraries. C.A.R., M.Me., Y.M., M.Mu., R.V., Ž.M., and N.A. developed the PAIR-Scan co-culture. C.A.R., M.Me., M.B., F.D., Ž.M., R.V., Y.W., developed PAIR-Scan screen sorting. M.Me., B.M., F.T.B.T.B., B.P.N., and J.U. developed single-well paired TCR- and neoAg-barcode-sequencing. J.W., Y.M., M.Mu., M.Me., C.A.R., R.V., Ž.M., B.O.S., and Y.W. performed and validated PAIR-Scan screens. M.Me., B.P.Y.K., and C.A.R. designed TCR and Ag libraries and wrote the PAIR-Scan screen analysis code. M. Me., T.N.S. and C.A.R., wrote the manuscript. T.N.S., W.S., and C.J.W. supervised the project. All authors reviewed and revised the manuscript.

## Ethics declarations

T.N.S. is an advisor to Allogene Therapeutics, Merus, Neogene Therapeutics, JNJ, and Scenic Biotech and is stockholder in Allogene Therapeutics, Cell Control, Celsius, Merus, Polar Therapeutics and Scenic Biotech. T.N.S. is also a venture partner at Third Rock Ventures, all outside the submitted work. J.B.A.G.H. is an advisor to AstraZeneca, BioNTech, BMS, Immatics, Iovance, Ipsen, Molecular Partners, Neogene Therapeutics, Novartis, Sastrā, T-Knife, Third Rock Ventures and Tzu; is a recipient of research grant support from Asher Bio, BioNTech, BMS and Sastrā; and is a stock option holder of Sastrā. C.J.W. is an equity holder of BioNTech; and is a scientific advisory board member of Repertoire, Aethon Therapeutics, Nature’s Toolbox and Adventris.

---

**Extended Data Fig. 1:**
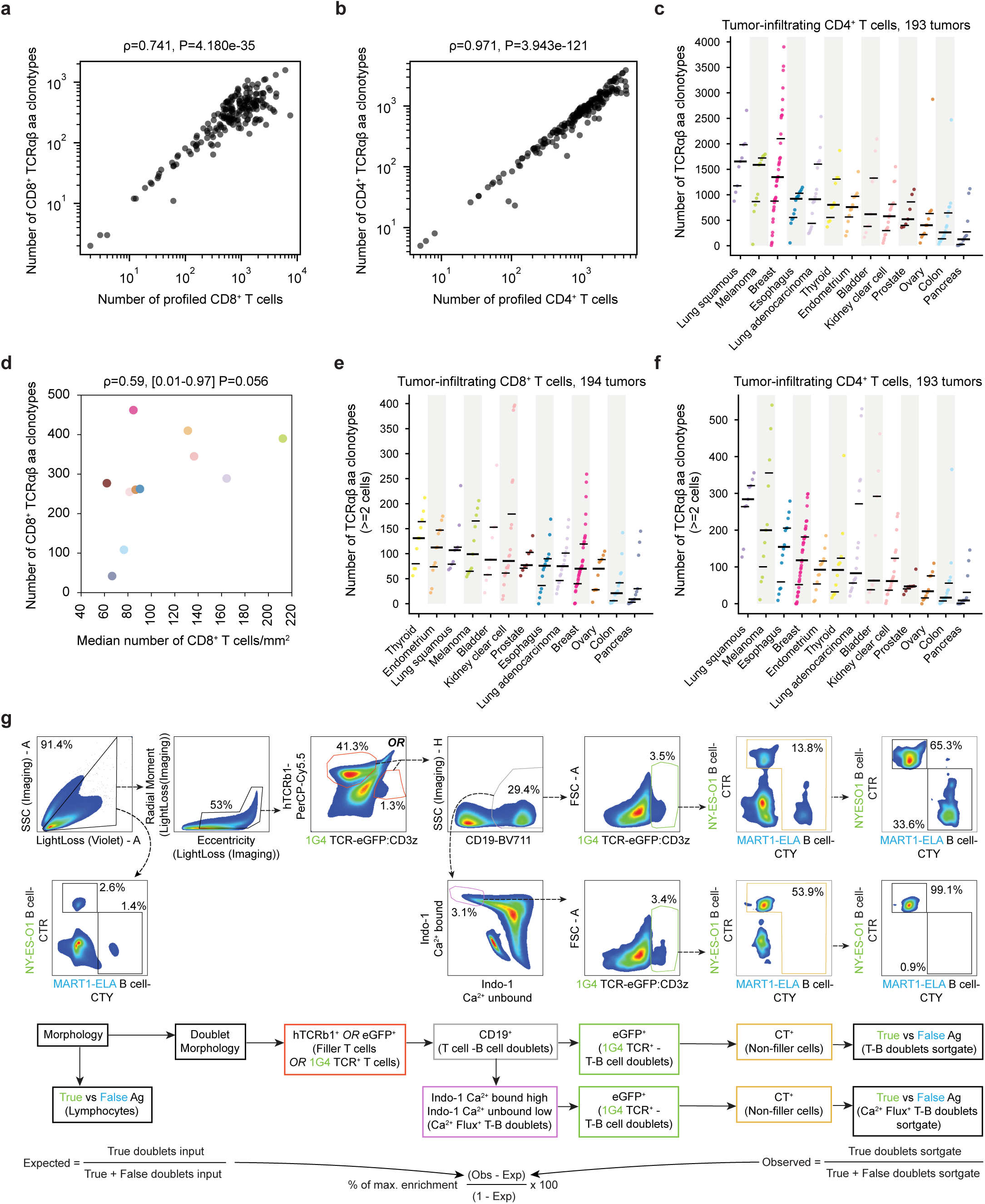
Number of TIL clonotypes across 13 cancer types and flow cytometric gating strategy for doublet quantification. **a, b)** Scatterplots depict correlations between total number of profiled T cells and detected number of unique clonotypes (ρ and P values indicated, two-sided Spearman correlation). Dots represent individual tumor samples. **c)** Number of unique tumor-infiltrating CD4^+^ T cell TCRαβ amino acid clonotypes identified in 193 tumors by single-cell sequencing across 13 cancer types (studies listed in Supplementary Table 1). Dots represent individual tumors. Lines indicate the median and the 25th/75th percentiles of the number of clonotypes. (**d**) Scatterplot depicting the correlation between the median number of intratumoral CD8^+^ T cells/mm² and the median number of intratumoral CD8^+^ T cell TCRαβ amino acid clonotypes (Fig. 1a) across 10 cancer types, colored as in (b) (two-sided Spearman’s ρ, 95% CI, and P value indicated). The number of intratumoral CD8^+^ T cells/mm² was quantified in 2,023 tumors using immunofluorescence staining and whole-slide imaging^14^. The cancer type “Non-Small Cell Lung” in^14^ was mapped to “lung adenocarcinoma” in Fig. 1a, as this represents the major NSCLC subtype. **e, f**) Number of clonally expanded (≥2 cells) tumor-infiltrating CD8^+^ (**e**) and CD4^+^ (**f**) amino acid clonotypes (studies listed in Supplementary Table 1). Dots represent individual tumor samples. Lines indicate the median and the 25th/75th percentiles of the number of clonotypes. **g)** Flow cytometry gating strategy and calculation to determine enrichment of true over false T–B cell doublets in simulated library-on-library co-cultures in Fig. 1e,g. Doublets are identified by morphology and marker expression ([hTCRb1^+^ *OR* 1G4 TCR^+^] *AND* CD19^+^). True and false conjugates are doublets identified as 1G4 TCR^+^ (eGFP^+^) doublets expressing cognate (NY-ESO-1, CellTrace Red, CTR) or control (MART1-ELA, CellTrace Yellow, CTY) antigen, respectively.

**Extended Data Fig. 2:**
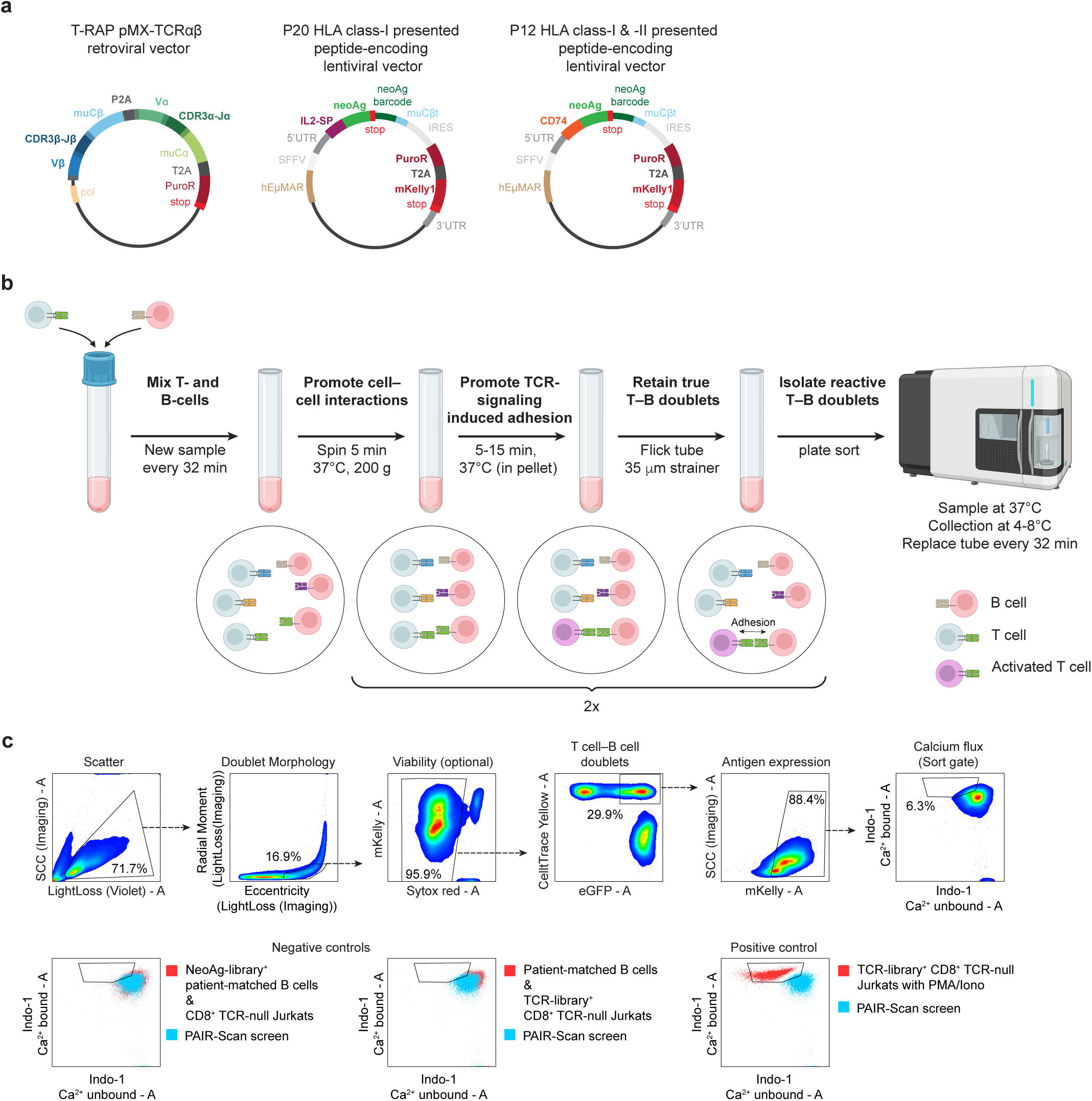
PAIR-Scan vector design and screening parameters. **a)** Schematic maps of PAIR-Scan expression vectors. T-RAP pMX-TCRαβ is a retroviral transfer plasmid for the expression of libraries of human TCRα and TCRβ chains (featuring mouse TCR constant domains) and a puromycin resistance gene, separated by 2A elements^21^. P20 and P12 are lentiviral transfer plasmids for peptide expression. P20 contains a 5′ IL-2-derived signal peptide leader sequence^19^, the neoAg or minimal peptide, and a stop codon. P12 contains a CD74 leader sequence to achieve trafficking to the endosomal compartment for MHC class II loading^12,65^. Each vector is designed to contain a neoantigen barcode and a portion of the murine constant TCRβ chain (non-coding, shared with T-RAP pMX-TCRαβ) that serves as a common reverse transcription priming site. Both P20 and P12 encode an internal ribosome entry site (IRES)-driven puromycin resistance gene product and the mKelly1 fluorescent protein, separated by 2A elements. In addition, both contain the enhancer (Eμ) and the matrix/scaffold-attachment regions (MARs) of the human immunoglobulin heavy chain which, combined with the spleen focus-forming virus (SFFV) promoter, boosts transgene expression in B cells^42^. **b)** PAIR-Scan workflow. Transduced T cell and B cell libraries are mixed and centrifuged (5 min, 200 g, 37 °C) to promote cell–cell contact^66^, followed by incubation of cell pellets (5–15 min, 37 °C) to allow TCR signaling–induced adhesion. Gentle tube flicking dissociates transient non-specific contacts while retaining stable cognate T–B cell doublets. Suspensions are passed through a 35 µm filter and sorted into 384-well collection plates, according to (c). As doublets eventually dissociate, and both adhesion and Ca^2+^ flux are temperature dependent^24,26,66^, sample tubes are maintained at 37 °C and replaced every 32 min. Plates are maintained at 4–8 °C until further processing. **c)** Gating hierarchy and experimental controls for PAIR-Scan screen sorts. Top row: Identification of doublets by scatter (Side Scatter vs Light Loss) and imaging morphology parameters (Radial Moment vs. Eccentricity), optional viability selection (Sytox red^-^), T–B doublet gating (CellTrace^+^eGFP^+^), target antigen expression (mKelly1^+^), and sorting of cell doublets displaying calcium flux (indo-1 Ca^2+^-bound^high^ /indo-1 Ca^2+^-unbound^low^, Ca^2+^-flux^+^). Bottom row: Overlay of calcium flux gate boundaries on negative controls (NeoAg-library^+^ autologous B cells co-cultured with CD8^+^ TCR-null Jurkat cells, and Ag-null patient–matched B cells co-cultured with TCR-library^+^ CD8^+^ TCR-null Jurkat cells) and positive control (TCR-library^+^ CD8^+^ TCR–null Jurkat cells treated with PMA/Ionomycin) relative to the PAIR–Scan screening signal.

**Extended Data Fig. 3:**
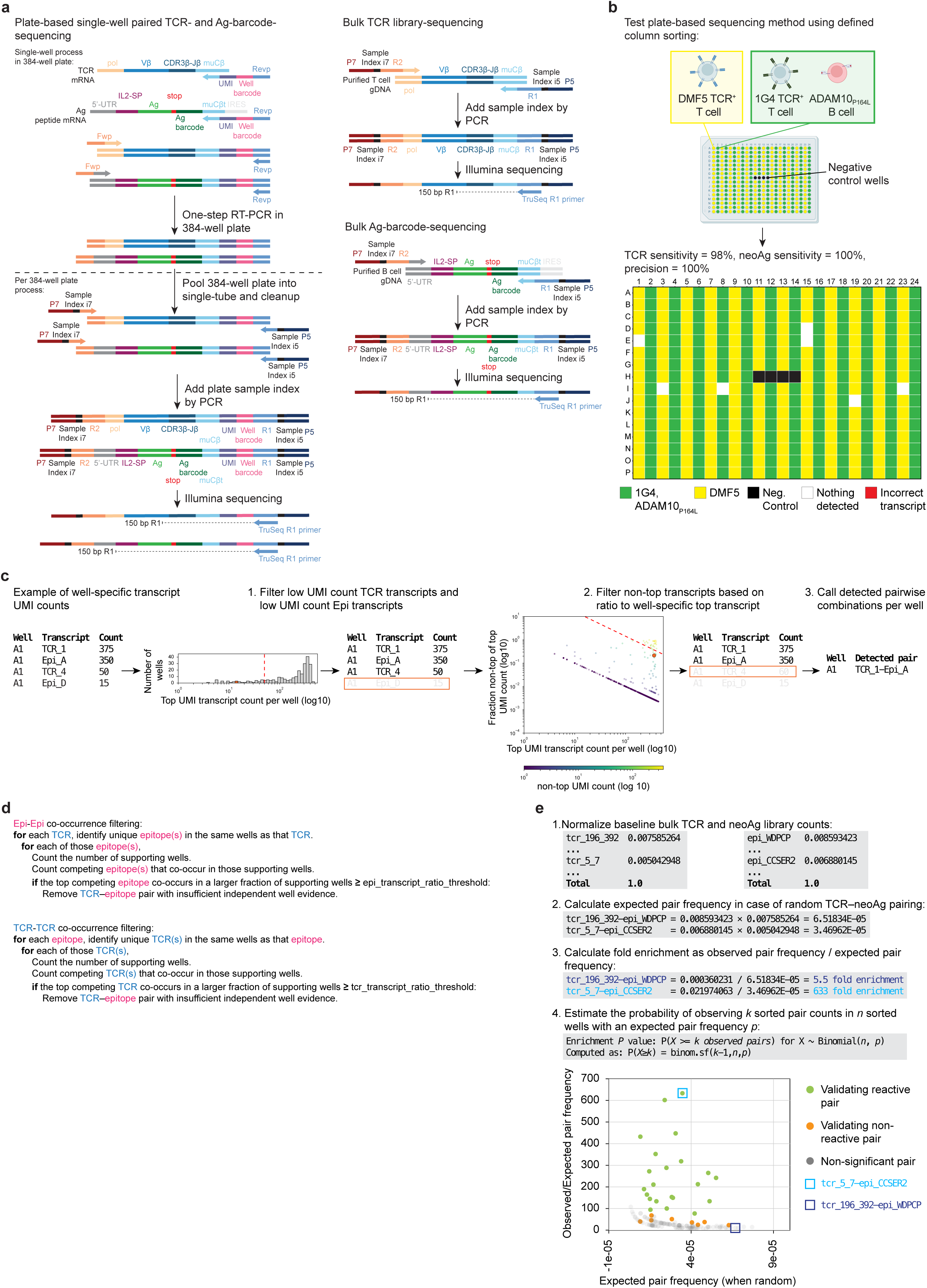
Single-well plate-based sequencing and data preprocessing. **a)** Single-well RNA sequencing schemes for paired TCR–antigen (Ag) detection in 384-well plates (left) and bulk DNA sequencing for TCR and Ag abundance quantification (right). Target mRNA from sorted doublets is used to perform a one-step RT-PCR with well-specific barcode tagging and unique molecular identifier (UMI) tagging. Wells are pooled and amplified using primers containing plate–specific unique dual indexes (UDIs) and Illumina adapters for subsequent deep sequencing. Genomic DNA isolated from TCR and neoantigen libraries is amplified using PCR primers targeting vector sequences and containing Illumina adapters for deep sequencing. **b)** Validation of single-well sequencing accuracy using a defined 384-well sorting experiment. Columns were seeded with defined T cell–B cell combinations (1G4-TCR^+^ Jurkat cells plus ADAM10_P164L_^+^ B cells, or DMF5-TCR^+^ Jurkat cells) or negative controls (empty wells). Data confirms high concordance between sorted input and sequencing output, achieving 98% TCR sensitivity, 100% neoantigen sensitivity, and 100% precision. **c)** Well-specific UMI filtering and TCR–Ag calling. Filtering removes low-abundance transcripts and secondary transcripts based on their UMI count ratio relative to the top transcript per well Filtering is separately performed for TCR and Ag transcripts. TCR–Ag pairs are then called per well as the pairwise combinations of the retained TCR and Ag transcripts. **d)** Epi-Epi and TCR-TCR co-occurrence filtering removes TCR–Ag pairs with insufficient independent supporting wells. For each TCR, a TCR–Ag pair is removed if a second Ag co-occurs in these well above a set threshold (e.g., 0.5 in this example) of its supporting wells, ensuring that only TCR–Ag pairs with sufficient independent well evidence are retained. TCR–TCR co-occurrence is filtered in the same manner. **e)** Calculation of PAIR-Scan confidence for individual TCR–Ag pairs and calculation of TCR–Ag pair screen fold enrichment. PAIR-Scan confidence is the probability of observing *k* sorted pair counts in *n* sorted wells given an expected random pairing frequency *p*. One highly confidence pair (tcr_5_7–epi_CCSER2) and one low confidence pair with high expected random pairing frequency (tcr_196_39–epi_WDPCP) are shown as illustrative examples.

**Extended Data Fig. 4:**
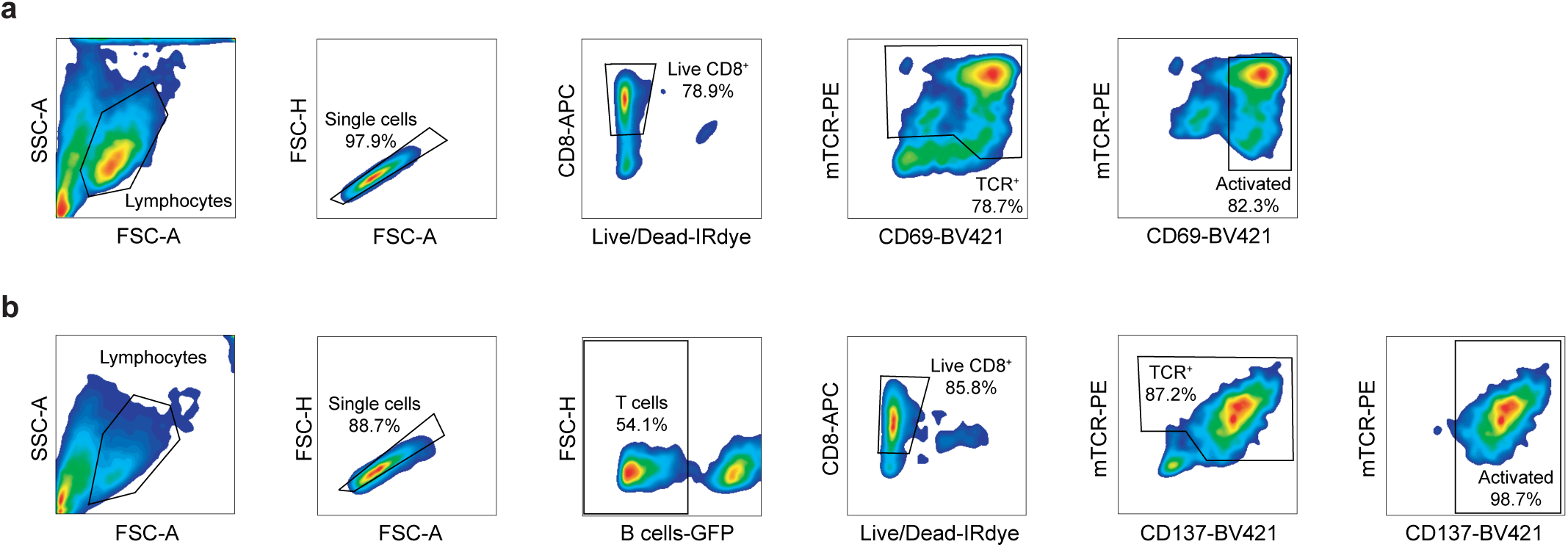
Gating strategy for arrayed TCR–neoAg pair validations. **a)** Gating strategy for validation of TCR–neoantigen (neoAg) pairs using Jurkat cells. CD8^+^ TCR-null Jurkat cells transduced with candidate TCRs were co-cultured with patient-derived autologous B cells expressing the indicated neoantigens. T cell reactivity was assessed by quantification of activated (CD69^high^), viable T cells (CD8^+^IRDye^-^TCR^+^). **b)** Gating strategy for validation of TCR–neoAg pairs using primary T cells. Primary CD8^+^ T cells transduced with candidate TCRs were co-cultured with patient–matched B cells (eGFP^+^) expressing indicated neoantigens. T cell reactivity was assessed by quantification of activated (CD137^high^), viable T cells (CD8^+^IRDye^-^eGFP^-^TCR^+^).

**Extended Data Fig. 5:**
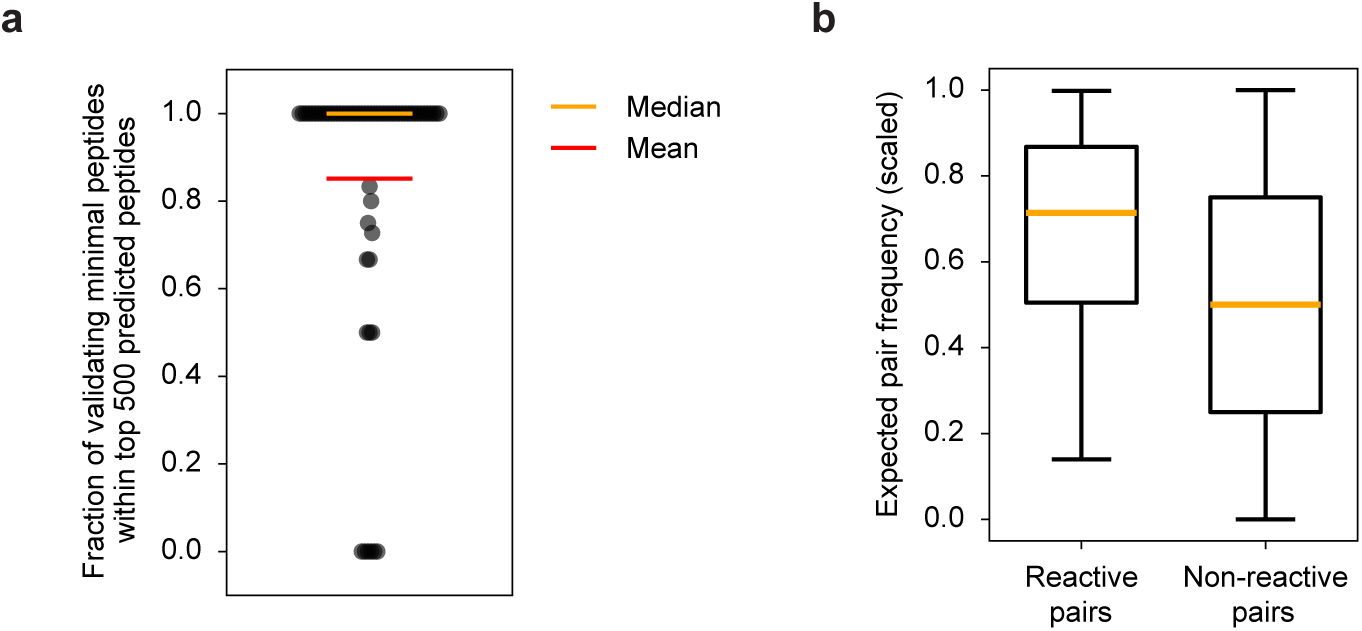
Additional validation of minimal peptide ranking and PAIR-Scan. **a)** Fraction of experimentally validating reactive minimal peptides included in the top 500 ranked minimal peptides per patient in the Gartner et al. dataset^31^. Reactivity of tumor-infiltrating lymphocytes (TIL) of 70 metastatic cancer patients was screened towards nonsynonymous expressed mutations (median 125 mutations per patient, range 1–679 mutations). For TIL-recognized neoAg(s) (median 1 recognized neoAg per patient, range 1–11 neoAgs), minimal peptides required for recognition were experimentally identified, giving the opportunity to test whether this minimal peptide would be included in a top 500 ranked minimal peptide set. Minimal peptides were ranked by predicted HLA binding confidence and mRNA expression. Each dot represents one patient; orange line, median (100%); red line, mean (85%). **b)** Expected pair frequency (percentile rank scaled) of reactive and non-reactive TCR–neoAg pairs across PAIR-Scan screens of patients A-D. Expected pair frequencies were calculated using bulk TCR and neoAg library sequencing data as depicted in Extended Data Fig. 3e. Boxes indicate median and 25th/75th percentiles of the expected pair frequencies; whiskers show values within 1.5× the interquartile range, with individual points indicating outliers beyond the whiskers.

**Extended Data Fig. 6:**
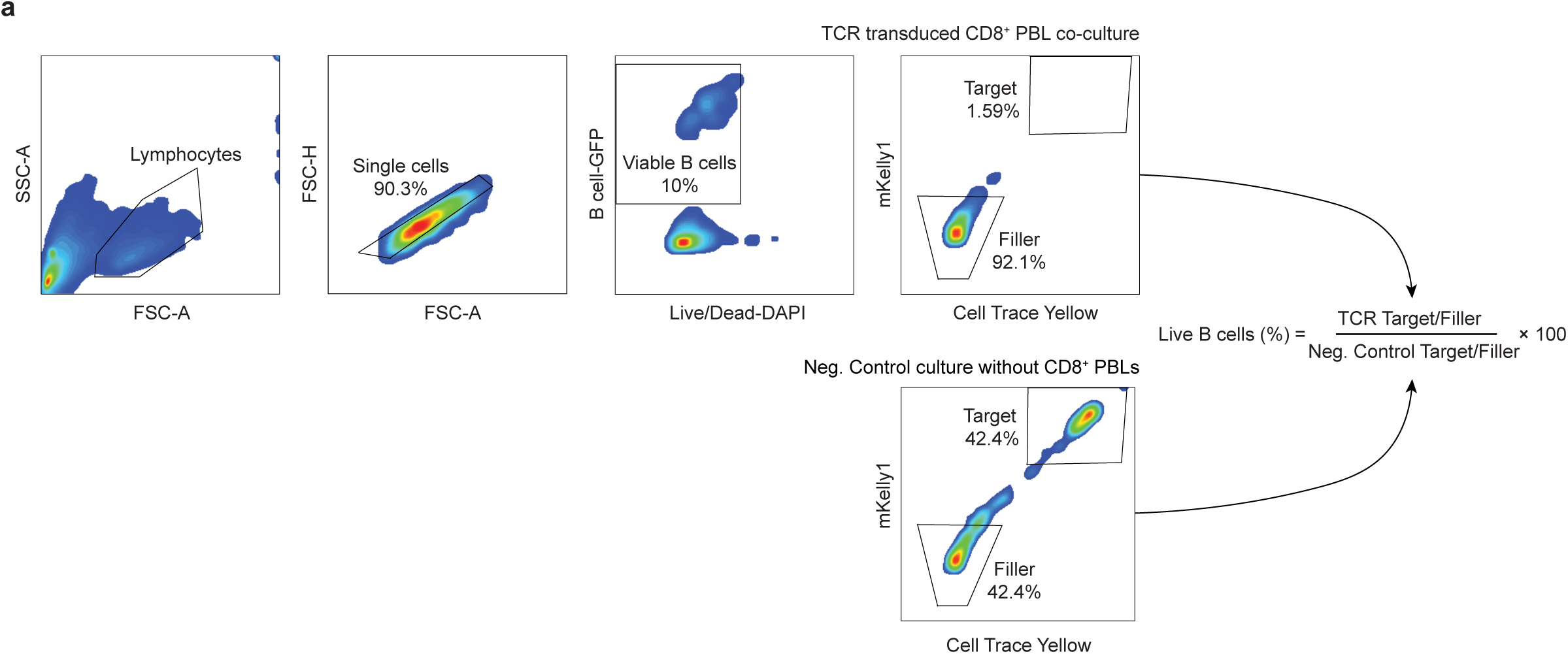
Gating strategy for cytotoxicity assays. **a**) Gating strategy and quantification of T cell-mediated killing of neoAg-expressing B cells. Primary CD8^+^ T cells transduced with candidate TCRs were co-cultured with neoAg^+^ target B cells (mKelly^+^ CellTrace Yellow^+^, eGFP^+^) and filler B cells (unmodified, eGFP^+^) pre-mixed at a 1:1 ratio. Antigen-specific killing was determined by analyzing the ratio of viable (DAPI^-^) target cells and filler cells in the presence or absence of T cells.

